# Recognition and Resolution of KRAS 5⍰UTR RNA G-Quadruplexes by hnRNPA1

**DOI:** 10.64898/2026.02.13.705690

**Authors:** Zahraa Othman, Matthieu Ranz, Ylenia Cortolezzis, Pedro Lourenço, David Moreira, Ahmad Daher, Carla Cruz, Eros Di Giorgio, Luigi E. Xodo, Gilmar F. Salgado

## Abstract

The KRAS oncogene, central to cellular signaling via MAPK and PI3K-AKT pathways, is a notorious cancer driver frequently activated in pancreatic, colorectal, and lung carcinomas. Regulation of human KRAS oncogene expression is important due to its capital role in cell growth, proliferation, and survival. Misregulation of its expression contributes directly to the development and progression of multiple types of cancer. In previous studies, the role of G-quadruplexes elements in both the promoter and 5’ UTR regions have shown to play important roles in *KRAS* expression, particularly when these G4s elements interact with regulatory protein hnRNPA1. In this study, we reveal that *KRAS* expression is also modulated at the post-transcriptional level through the formation of RNA G-quadruplexes (rG4s) situated at the 5⍰ untranslated region (5⍰UTR) of the mRNA. Biophysical and binding studies were carried out to probe the interaction. Through isothermal titration calorimetry (ITC), we quantified a strong binding affinity between the UP1 domain of hnRNPA1 and short-nucleotide RNA segments capable of adopting different G-quadruplex fold. The binding interaction is characterized by a favorable Gibbs free energy change in the range of ΔG ≈ -32 to -34 kJ/mol, suggesting a specific and energetically favorable association. One-dimensional and two-dimensional ^1^H-^15^N HSQC NMR spectroscopy revealed pronounced chemical shift changes in residues of both RNA recognition motifs (RRMs) of UP1, signifying direct contact with the rG4 structure.

## Introduction

Heterogeneous nuclear ribonucleoprotein A1 (hnRNPA1) was among the earliest hnRNPs identified in the 1980s, emerging from pioneering biochemical fractionation and immunological techniques that revealed nuclear proteins tightly associated with pre-mRNA (1-4). Initial studies, conducted in HeLa and hepatoma cells, demonstrated that removal of protein components from hnRNP-mRNA complexes impaired translation, suggesting an essential role for these proteins in translation factor recruitment and translational regulation (5,6). Among the hnRNP family members, hnRNPA1 emerged as one of the most abundant and characterized proteins. Early work determined its molecular weight to be approximately 34⍰kDa (7), described its nucleocytoplasmic shuttling behavior, and characterized its high-affinity, single-stranded RNA binding mediated by conserved RNA recognition motifs (RRMs) (8). Initial interest focused on its roles in pre-mRNA splicing, nuclear export, and mRNA stability key processes in gene expression and regulation, positioning hnRNPA1 as a model for RNA-binding protein function (9). Encoded by the HNRNPA1 gene, hnRNPA1 exists in two primary isoforms: hnRNP⍰A1 (A1-A), the canonical 320-amino acid (≈341⍰Da) form, and hnRNP⍰A1B (A1-B), which includes an additional exon (7B), resulting in a 372-amino acid (≈381⍰Da) protein (10). hnRNPA1 is abnormally expressed at different stages of cancer progression. Its cellular levels are correlated with pathophysiological phenotypes and clinical prognosis of cancers, indicating hnRNPA1 may be an important cancer biomarker and therapeutically target (11). Structurally, the protein comprises two major domains: (i) an N-terminal region containing tandem RRMs (residues⍰1-196, known as UP1), and (ii) a C-terminal glycine-rich domain (GRD) featuring low sequence complexity, an RGG-box, and the M9 nuclear localization/export signal (12). The tandem RRMs (RRM1 and RRM2) adopt a conserved βαβββα fold, with RNP2 and RNP1 motifs on β-strands 1 and 3 mediating RNA binding specificity (13-15). High-resolution structures of UP1 complexed with RNA or telomeric DNA have revealed cooperative domain interactions and the importance of inter-RRM linkers; notably, conserved salt bridges (R75:D155 and R88:D157) stabilize a “lid” bridging the two RRMs (16,17). In contrast, the C-terminal domain is intrinsically disordered, facilitating interactions with proteins and RNA, enabling nucleocytoplasmic transport, and driving phase separation. It also contains an RGG-box that mediates nucleic acid binding and phase behavior, as well as the M9 motif for transportin - mediated nuclear import/export (12,18). Multiple studies have demonstrated that this low-complexity region promotes liquid-liquid phase separation (LLPS) and formation of stress granules and amyloid-like fibrils (19-22). Functional analyses have shown non-redundant contributions of each RRM: mutagenesis and domain-swap experiments revealed that disruption of one RRM impairs hnRNPA1’s ability to regulate splicing and RNA transport (23). The RGG domain further enhances binding to guanine-rich RNA via cation-π interactions, and its arginine residues are methylated by PRMTs, modulating both RNA binding and LLPS propensity (24-26). Advanced structural studies, including cryo-EM, NMR, SAXS, cross-linking mass spectrometry (XL-MS), and single-molecule FRET, have mapped the dynamic architecture of full-length hnRNPA1. These methods reveal a modular protein in which the RRMs form a compact core, while the C-terminal IDR remains flexible and engages in transient intramolecular contacts, likely serving an autoinhibitory role (27-29). Cryo-EM structures have elucidated the fibrillar architecture of the C-terminal domain, underscoring the roles of the PY-NLS and RGG motifs in amyloid-like filament formation (30,31). Post-translational modifications (PTMs) further regulate hnRNPA1 behavior: phosphorylation of serine residues (e.g., Ser192, Ser199) within the GRD introduces negative charges that weaken LLPS and RNA cooperation, whereas arginine methylation in the RGG box alters specificity and interaction profiles (26,33). Recent discoveries have expanded hnRNPA1’s functional repertoire beyond RNA metabolism, revealing its involvement in chromatin dynamics and transcriptional control via interaction with non-canonical nucleic acid structures. One prominent example involves the human KRAS gene, a central oncogenic driver frequently mutated or dysregulated in multiple cancers, particularly pancreatic ductal adenocarcinoma (PDAC). In addition to its well-characterized mutational profile, KRAS expression is subject to intricate layers of transcriptional and post-transcriptional regulation (34, 35). Of particular interest are guanine-rich (G-rich) sequences located in the KRAS promoter and 51 untranslated region (5⍰UTR), which can fold into G-quadruplex (G4) DNA or RNA structures respectively. These elements act as epigenetic-like switches that modulate transcriptional output and translation efficiency. Moreover, G4-binding specificity is modulated by the RGG-box domain of hnRNPA1 (36), which likely enhances recognition and unfolding efficiency. The KRAS-ILK-hnRNPA1 axis, as described by Chu and colleagues (37), was shown to function as a redox-sensitive feedback loop: oxidative stress promotes the recruitment of hnRNPA1, PARP1, and Ku70 to oxidized G4s (e.g., those harboring 8-oxoguanine) (37), thereby stimulating KRAS transcription under oncogenic stress conditions. In parallel, RNA G-quadruplexes (rG4s) in the KRAS 5⍰UTR have emerged as critical post-transcriptional regulators, impacting mRNA stability and translational control. Recent work (38) identified three conserved rG4s UTR-1, UTR-C, and UTR-Z that adopt parallel G4 topologies and reduce KRAS mRNA half-life. These structures act as cis-elements that hinder ribosomal scanning, repressing translation in a potassium-dependent manner (37, 38). Moreover, our recent data demonstrated that G4 formation at the 5⍰ UTR of KRAS preserves transcript stability by preventing antisense RNA binding. In this context, hnRNPA1 resolved the G4 structure by regulating the interaction between the KRAS 5⍰ UTR and antisense RNA (38). Taken together, the growing body of evidence positions hnRNPA1 as a central player in the multi-level regulation of oncogenic KRAS. Through its dual capacity to bind and remodel G4 structures in both DNA and RNA contexts, hnRNPA1 integrates transcriptional and translational control. In this work, we attempt to decipher the recognition mechanism on how UP1 would recognize and unfold such stable structures such as RNA G-quadruplexes found in the 5’ UTR region of KRAS mRNA.

## Results and Discussion

### CD and 1D NMR spectroscopy of 5⍰ UTR G-Quadruplex

Unless otherwise stated, all experiments were carried out with UP1 **(Figure 1)**. The three oligonucleotides were analyzed by CD spectroscopy and 1D NMR **(Figure 2)**. They exhibit a typical CD profile of a parallel topology, characterized by a positive peak with a maximum at ≈260 nm and a negative minimum at ≈240 nm. All three UTRs exhibit this spectral signature, but with varying ellipticity intensities, suggesting differences in folding efficiency, G4 population, or G4 topology complexity. In addition, the CD topology signature for the three oligos show a positive peak at ∼290 nm, indicative of structural heterogeneity or mixed G4 topologies, such as a minor extra contribution from hybrid conformation or antiparallel, which is small for UTR-1, average for UTR-C and important for UTR-Z. The extra shoulder at 290 nm is an important hallmark of significant conformational heterogeneity for UTR-Z but, nevertheless its Δ*T*_*m*_ is higher due to extra molecular size, including 2 extra guanine tracks. It is also possible to think that adjacent G4s can stack or stabilize one another through cooperative folding. In addition, one-dimensional (1D) proton (^1^H) NMR spectroscopy was used to probe the imino-pattern resonances (≈10-12 ppm), which report on Hoogsteen hydrogen bonding within G-quadruplex (G4) tetrads. All three sequences display significant imino intensity, consistent with a substantial fraction of molecules adopting G4 structures under the tested ionic conditions. UTR-1 and UTR-Z exhibit broad, highly overlapped imino peaks with strong overall signal intensity but limited peak resolution. This pattern suggests the presence of multiple conformational or oligomeric states that interconvert on an intermediate NMR timescale, a behavior previously reported for RNA G4s with short G-tracts or sequence interruptions that promote structural heterogeneity. Upon increasing the temperature to 46 °C, the UTR-1 spectrum becomes partially resolved, indicating thermal destabilization of minor or less stable conformers and a shift toward a simpler population. In contrast, UTR-Z shows minimal sharpening at elevated temperature, consistent with a more persistent ensemble of slowly exchanging conformations or higher-order stacked species, a feature that has been described for other highly stable RNA G4s that readily form multimers. In clear contrast to these two heterogeneous systems, UTR-C exhibits multiple sharp imino resonances and well-dispersed aromatic peaks even at 25 °C, indicative of a predominantly monomeric and well-defined G4 topology. The presence of several fully resolved imino resonances is characteristic of RNA G4s with limited conformational exchange and well-organized loop geometries. Together, these data reveal marked folding differences among the KRAS 5 ⍰ -UTR sequences, placing UTR-C in the category of more canonical, structurally homogeneous RNA G-quadruplexes, whereas UTR-1 and UTR-Z populate broader structural ensembles reminiscent of previously reported non-canonical or multi-state RNA G4 systems. To resume, the broad line shapes of UTR-1 and UTR-Z are also compatible with transient end-to-end stacking or higher-order G4 oligomerization from NMR sample conditions, whereas UTR-C is mostly monomeric even at high concentrations.

**Figure 1.**
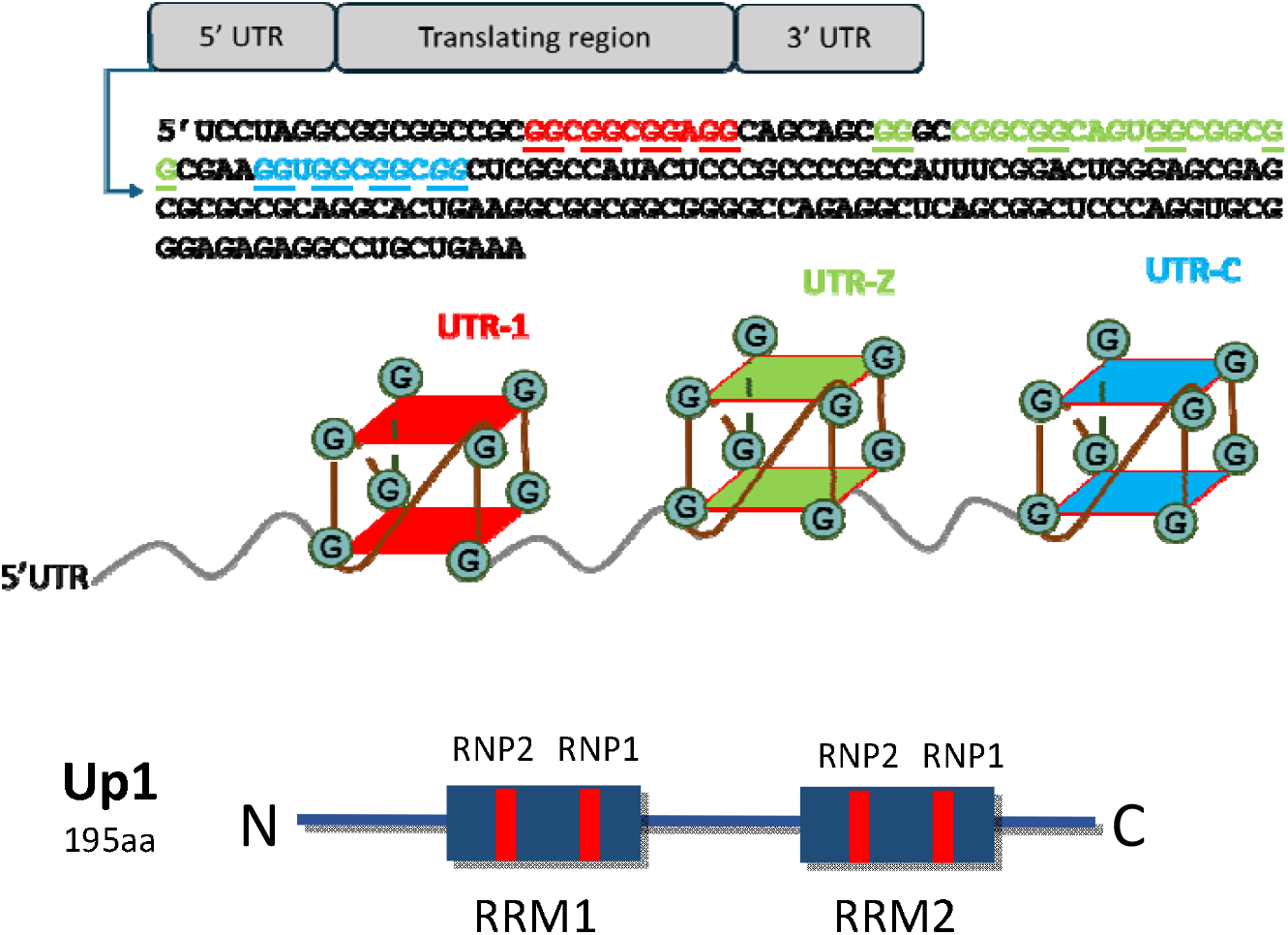
Sequence of KRAS 5’UTR region and UP1 domains. KRAS 5’UTR region contains three stretches of guanines that autoassemble into G-quadruplexes structures *in vitro*, they are color-coded as UTR-1, UTR-Z and UTR-C. In total we can observe 33 pairs of Gs. The UP1 domain organization is shown, including the conserved RNP2 and RNP1 sub-motifs within each RRM domain. In RRM1, the RNP2 and RNP1 motifs contain the conserved redidues KLFIG and RGFGF respectively. In RRM2, RNP2 and RNP1 contain the sequences KIFVG and RGFAF respectively.

**Figure 2.**
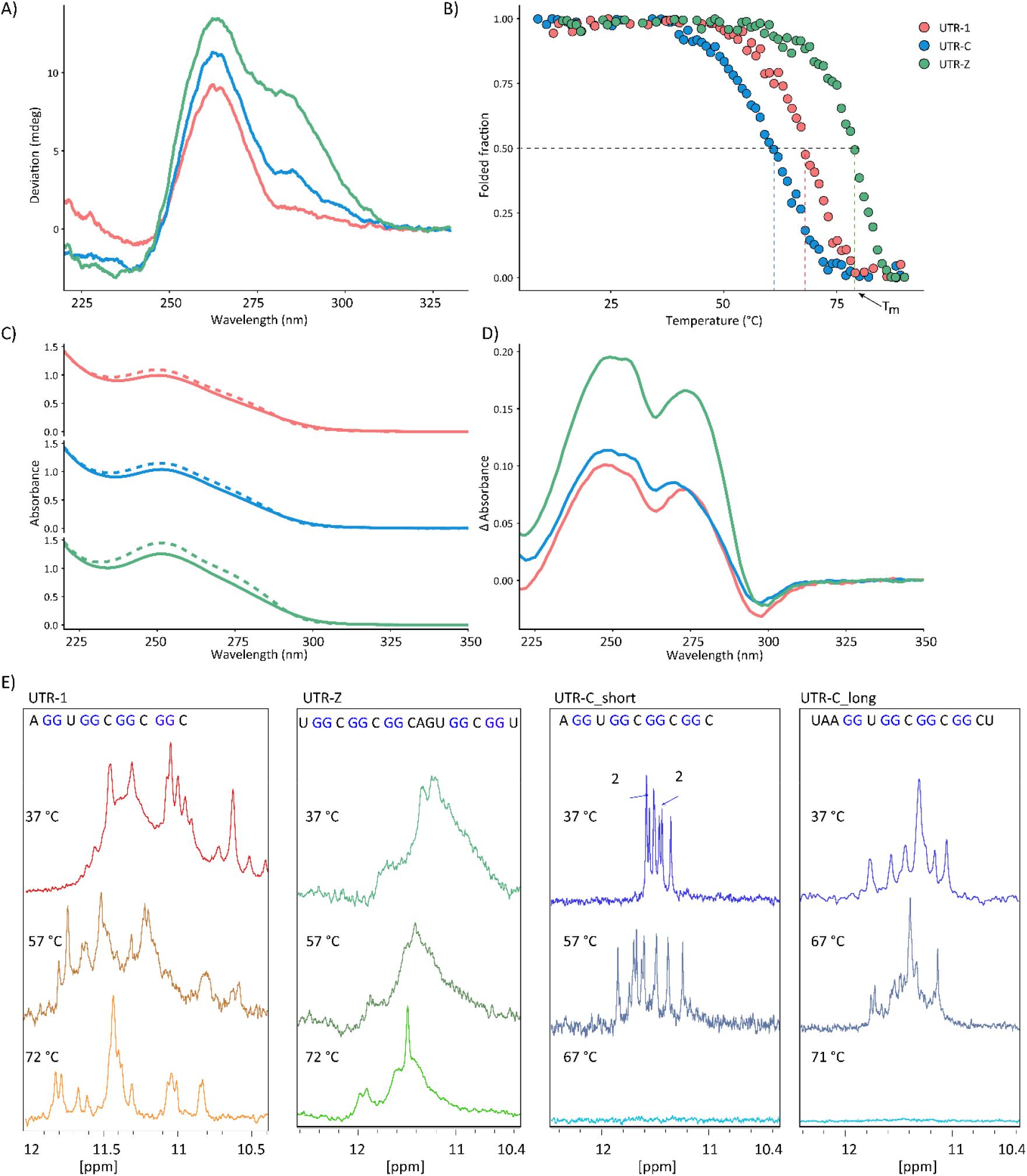
Biophysical characterization of KRAS 5UTR RNA G-quadruplexes. **(A)** Far-UV CD spectra of UTR-1 (red), UTR-C (blue), and UTR-Z (teal) recorded at 22 °C (3 µM RNA) in 10 mM KPi containing 50 mM KCl. All sequences display a dominant positive band centered at ≈260-265 nm and a negative band near ≈240 nm, consistent with parallel-type G-quadruplex folding. Differences in ellipticity intensity and band shape reflect sequence-dependent stacking interactions and conformational homogeneity. Spectra were baseline-corrected and processed identically for comparison. **(B)** Thermal unfolding profiles monitored by CD and plotted as folded fraction versus temperature. Folded fractions were normalized using pretransition and post-transition baselines for each construct. The sigmoidal transitions indicate cooperative unfolding, with UTR-Z exhibiting the highest apparent thermal stability, followed by UTR-1 and UTR-C. Apparent melting temperatures (T_m_), determined from the first derivative of the melting curves, are approximately 61 °C (UTR-C), 69 °C (UTR-1), and 80 °C (UTR-Z). **(C)** UV absorbance spectra acquired at 10 °C (solid lines) and 80 °C (dashed lines), illustrating temperature-dependent spectral changes associated with quadruplex unfolding. **(D)** Thermal difference spectra (TDS) obtained by subtracting the 10 °C spectrum from the 80 °C spectrum. Characteristic positive features in the 240-280 nm region are consistent with G-quadruplex signatures and enable comparison of relative base stacking contributions among the three motifs. **(E)** Sequences of the rG4 motifs located within the first 80 nucleotides of the KRAS 5⍰UTR and representative 1D-H NMR imino proton spectra (≈ 10-12.5 ppm) of UTR-1, UTR-Z, UTR-C_short, and UTR-C_long recorded at increasing temperatures (37-72 °C) in 50 mM KCl, 10 mM potassium phosphate buffer. Well-resolved imino resonances at lower temperatures confirm Hoogsteen hydrogen-bonded guanine tetrads, whereas progressive signal broadening and disappearance at elevated temperatures reflect thermal destabilization and unfolding. Sharper and more persistent imino signals observed for UTR-C_short indicate increased conformational homogeneity relative to the longer constructs.

### hnRNPA1 selectively binds rG4-containing regions within the KRAS 5⍰UTR in cells

To identify the regions within the KRAS 5⍰UTR predominantly bound by hnRNPA1, we generated three lentiviral reporters containing mScarlet fused to either the control sequence or individual UTR fragments (UTR -1, UTR -C, and UTR -Z). These lentiviral vectors were used to infect Panc-1 cells, either wild-type or hnRNPA1 knockout (**Figure 3**). Subsequently, RNA immunoprecipitation was performed using anti-hnRNPA1 or IgG antibodies, followed by RT-qPCR amplification of mScarlet. The data demonstrate preferential binding of hnRNPA1 to UTR - Z and UTR -C, consistent with the higher propensity of these regions to form rG4 structures in vivo (38). Overall, these findings highlight the KRAS 511UTR, particularly UTR -C and UTR -Z as strong binding sites for hnRNPA1.

**Figure 3.**
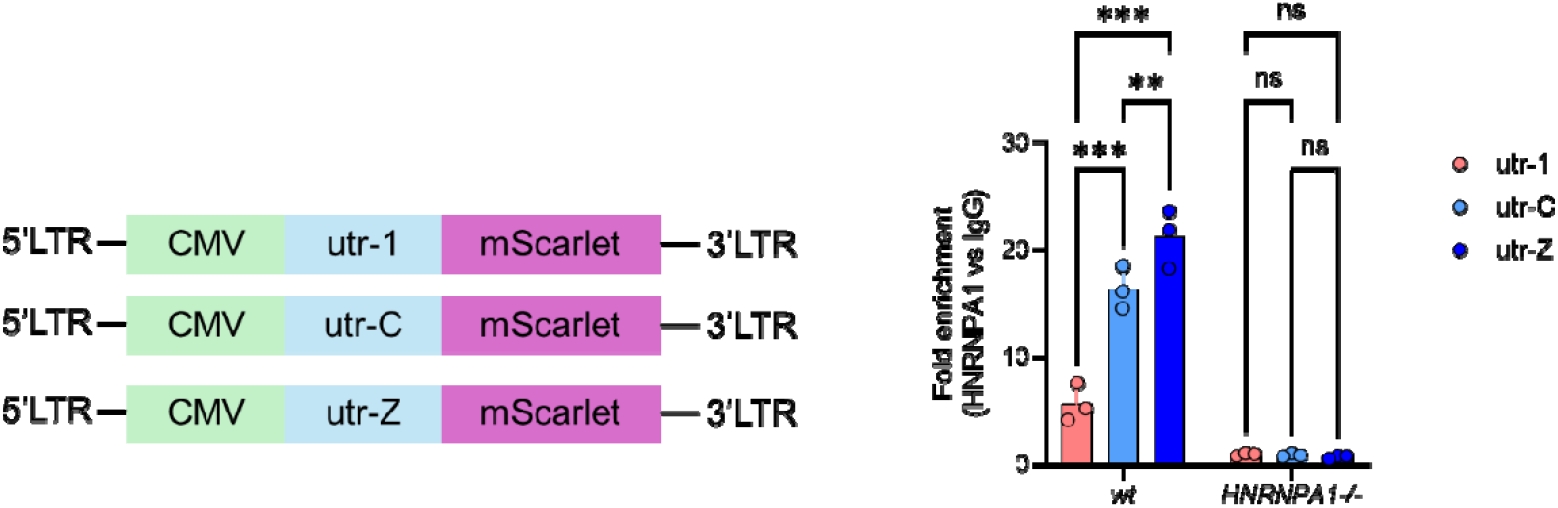
hnRNPA1 associates with KRAS 5⍰UTR reporter transcripts in cells. **(Left)** Schematic representation of lentiviral reporter constructs in which the mScarlet coding sequence is expressed under the control of the CMV promoter and preceded by the KRAS 5lllUTR containing either the UTR-1, UTR-C, or UTR-Z rG4 motif. Lentiviral particles were used to stably transduce wild-type (WT) or hnRNPA1 knockout (hnRNPA1^−/−) Panc-1 cells, generating cellular reporters to interrogate rG4-dependent RNA-protein interactions in vivo. **(Right)** RNA immunoprecipitation (RIP) assay performed using 3 µg of anti-hnRNPA1 antibody or isotype-matched IgG control. Immunopurified RNAs were reverse-transcribed and quantified by qPCR using primers targeting the 5⍰ region of the mScarlet transcript. Enrichment is reported as fold change relative to IgG control for each construct. Data represent mean ± standard deviation from three independent biological replicates (n = 3). Individual data points are shown. Preferential enrichment of KRAS 5⍰UTR reporters in hnRNPA1 immunoprecipitates relative to IgG supports selective association of hnRNPA1 with rG4-containing transcripts in cells.

### Thermodynamic Analysis of UP1 Binding to 5⍰ UTR G-Quadruplex RNAs by ITC

ITC experiments were performed both with and without the presence of soluble partner of UP1 Glutathione S-transferase (GST). GST is known to be an excellent partner that facilitates solubilization of some complex that may have some precipitation issues during titrations. **Figure 4** presents ITC measurements of the interaction between the UP1 protein (UP1 is a proteolytic fragment of hnRNPA1 that maintains binding to G4 RNA and G4 DNA and capacity unfold G4 structures), and three distinct RNA G-quadruplex-forming the 5⍰ untranslated region (UTR) constructs: UTR-C, UTR-Z, and UTR-1. The top panels show raw heat changes over time (μcal/sec) upon successive UP1 injections, while the bottom panels display integrated binding isotherms fitted to a single-site binding model, revealing the thermodynamic parameters of interaction. All three RNAs bind UP1 in the low or submicromolar range, with UTR-C showing the highest affinity (*K*_D_ ≈ 0.621μM). The ΔG values are relatively similar across constructs (−31 to −36⍰kJ/mol), suggesting that overall binding is thermodynamically favorable and mostly enthalpically driven. Enthalpy contributions (ΔH) are highest (most negative) for UTR-Z (−39.6 kJ/mol), suggesting strong direct favorable molecular interactions indicating more molecular contacts such as hydrogen bonding and stacking interactions. The lower ΔH for UTR-1 and-C (near −36 kJ/mol) may reflect either less optimal contacts or more conformational adaptation upon binding from a shorter nucleotide chain. In addition, UTR-Z pays a higher entropic cost, which is consistent with its strong 290 nm shoulder in CD, a hallmark of conformational heterogeneity. UP1 binds G4 RNA through enthalpically driven mechanisms, especially in UTR-Z, where strong interactions are countered by conformational locking. UTR-C, although slightly less enthalpically favorable, gains an entropic advantage, possibly reflecting minimal structural reorganization. These findings confirm the sensitivity of UP1 to RNA G4 conformational landscapes, and the need to consider both structure and dynamics in understanding RNA recognition by UP1.

**Figure 4.**
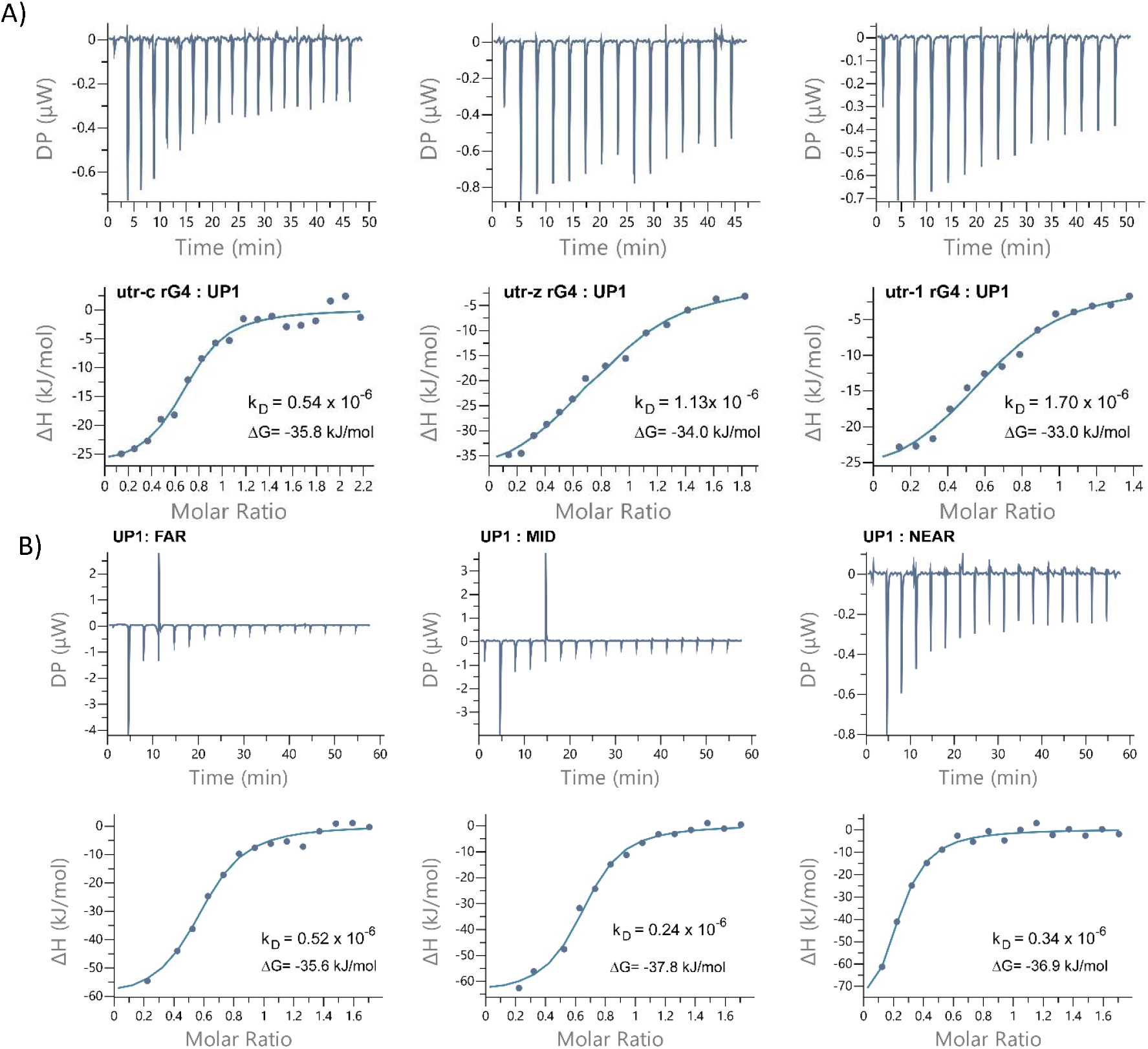
Thermodynamic characterization of UP1 binding to KRAS 5UTR rG4s and promoter G4 motifs by isothermal titration calorimetry (ITC). (A) Representative ITC raw thermograms (upper panels) and integrated binding isotherms (lower panels) obtained by titrating UP1 into solutions of UTR-C, UTR-Z, and UTR-1 rG4s. Solid lines correspond to global fits to a single-site binding model. Binding is exothermic for all three substrates and yields micromolar dissociation constants (**K**_*D*_) and favorable binding free energies (ΔG), as indicated in each panel. Differences in binding affinity and enthalpic contribution reflect sequence and topology dependent recognition of rG4 scaffolds by UP1. (B) Analogous ITC experiments performed using promoter-derived G4 sequences corresponding to FAR, MID, and NEAR motifs. Raw thermograms and fitted isotherms are shown with solid line fits to a single-site model. These constructs also exhibit exothermic binding with micromolar affinities, demonstrating that UP1 engages diverse G-quadruplex substrates using a conserved thermodynamic binding mode. For all experiments, heats of dilution were subtracted and data were analyzed using nonlinear least-squares fitting. Reported thermodynamic parameters (*K*_*D*_, ΔH, ΔG, and stoichiometry) are summarized in Table 1.

Concerning the promoter DNA G4s (far, mid and near): the K_*D*_ values (0.24-0.52 µM) and ΔG (≈ −36 to −38 kJ/mol) are in the same order of magnitude with previously reported values in the literature (39), 0.49-1.1 µM; ΔG −35.6 to −37.2 kJ/mol) for the near G4, since those types of results were never reported for the mid and far G4s. Concerning the 5⍰UTR RNA G4s (KRAS): the results obtained in this study (0.62-1.74 µM) are in the same order of magnitude as results found on studies with UP1-TERRA RNA G4 (≈1.7-2.0 µM, Ghosh & Singh 2020). RNA affinities appear slightly weaker than DNA, but there are not many such examples of G4s interacting with UP1 in order to have a broader perspective. Overall, our binding studies by ITC show a slighter better affinity of UP1 towards DNA than RNA G4. ITC data aligns well with published studies, supporting UP1’s role in KRAS recognition of both the promoter and UTR G4s.

**Table 1.**
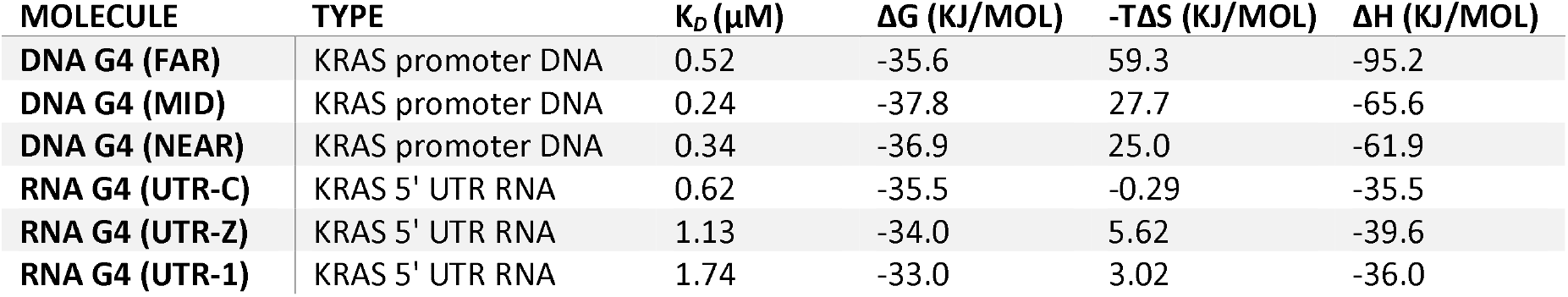
Resume of Thermodynamic parameters obtained from ITC experiments.

| MOLECULE | TYPE | $K_D$ (μM) | $\Delta G$ (KJ/MOL) | $-T\Delta S$ (KJ/MOL) | $\Delta H$ (KJ/MOL) |
| --- | --- | --- | --- | --- | --- |
| **DNA G4 (FAR)** | KRAS promoter DNA | 0.52 | -35.6 | 59.3 | -95.2 |
| **DNA G4 (MID)** | KRAS promoter DNA | 0.24 | -37.8 | 27.7 | -65.6 |
| **DNA G4 (NEAR)** | KRAS promoter DNA | 0.34 | -36.9 | 25.0 | -61.9 |
| **RNA G4 (UTR-C)** | KRAS 5' UTR RNA | 0.62 | -35.5 | -0.29 | -35.5 |
| **RNA G4 (UTR-Z)** | KRAS 5' UTR RNA | 1.13 | -34.0 | 5.62 | -39.6 |
| **RNA G4 (UTR-1)** | KRAS 5' UTR RNA | 1.74 | -33.0 | 3.02 | -36.0 |

### UP1 binding to KRAS 5⍰UTR rG4 measured by microscale thermophoresis (MST)

Accordingly with MST traces (Figure 5), the 3 oligonucleotides show strong and consistent fluorescence decay, indicative of a large conformational or hydration change upon binding. All three MST curves are clean and sigmoidal, indicating cooperative and specific interaction. UTR-Z displays the strongest affinity (K_*D*_ = 0.125⍰μM) for UP1. The MST results support the CD results showing mixed G4 populations (290 nm shoulder), and also the ITC isothermals showing strong enthalpic interaction but entropic cost. This evidence likely reflects UP1 recognition of flexible, heterogeneous G4 states, possibly due to conformational selection. The more intermediate binding observed for UTR-C (K_*D*_ = 0.215⍰μM), is consistent with a well-folded G4 structure with moderate plasticity. CD and ITC suggest this is a stable, parallel G4, and the MST confirms high binding efficiency despite lower flexibility. This likely reflects the binding of a canonical G4 scaffold, which is well-suited for RRM recognition but smaller in size compared to UTR-Z, which contains more nucleobases, longer loops, and a more polymorphic ensemble. UTR-1 has the weakest interaction (K_*D*_ = 7.67⍰μM), more than an order of magnitude weaker than UTR-Z. The respective MST traces are shallower, showing little thermophoretic shift, possibly due to minimal conformational change or partial folding of the G4. It matches the CD data with a low 265⍰nm/295⍰nm ratio (indicative of a less stable G4 folding), and the lowest *K*_D_ and less favorable enthalpy and entropy seen by ITC. Likely reflects insufficient G4 structure formation, or a sequence that does not present the correct topology for UP1 engagement. Strong thermophoretic shifts correlate with conformational rearrangement, consistent with UP1 selecting among multiple G4 topologies (firstly the UTR-Z then UTR-C and finally UTR-1). The weaker stability of UTR-1 compared to UTR-Z and UTR-C is in keeping with the finding that under physiological conditions two G4 RNA structures (that should be UTR-Z and UTR-C) within the same 5’UTR KRAS mRNA are stably folded, while the third (UTR-1) most probably exist in equilibrium between the folded and unfolded states (38). Together with CD and ITC, MST reveals a combined thermodynamic structural and binding set of data, where structural plasticity and interaction enthalpy drive UP1 recognition and it also indicates that UP1 has a clear binding preference for structurally heterogeneous or plastic G4s.

**Figure 5.**
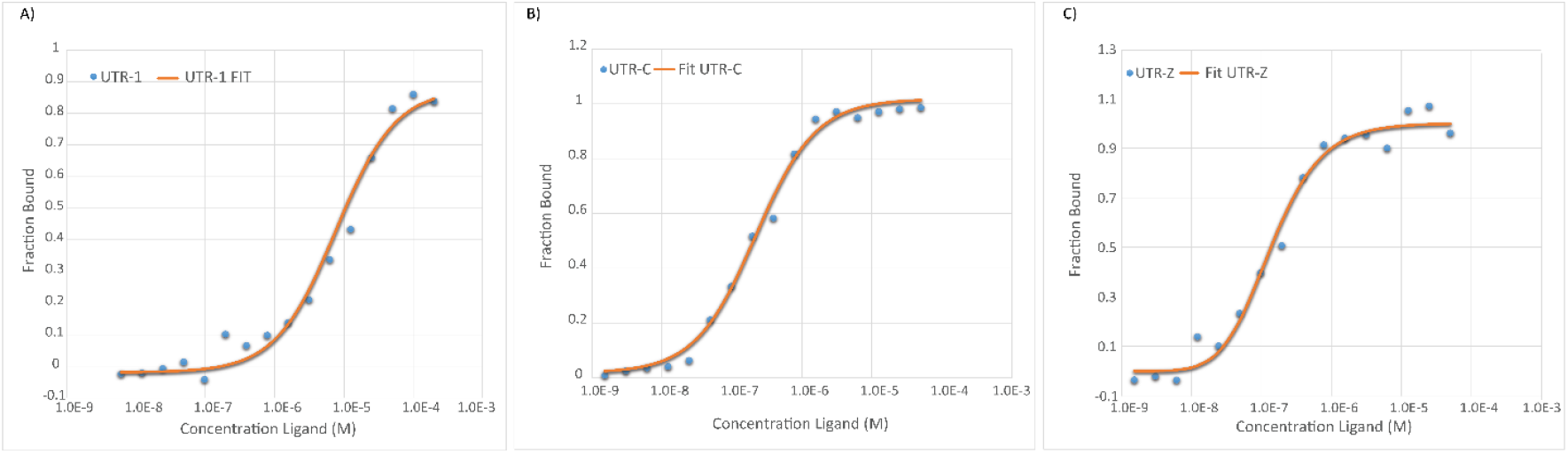
Microscale thermophoresis (MST) analysis of UP1 binding to KRAS 5UTR rG4 constructs (A-C). MST dose-response curves and fits (orange lines) for Cy5-labeled UTR-1 (A), UTR-C (B), and UTR-Z (C) (5 nM RNA, 1× KPi + 100 mM KCl + 0.05% Tween-20). UTR-Z shows the highest apparent affinity (KD = 0.125 µM), likely due to conformational selection from its heterogeneous ensemble; UTR-C binds robustly to its canonical scaffold. The resulting dose-response curves were fitted to a one-site binding model to extract K d values.

### Binding affinity measured by Fluorescence Titration

Fluorescence titration was also used to inspect the binding interaction at nano molar range between UP1 and pre-folded UTRs G4s double-labelled (FAM and TAMRA at the 5’ and 3’ respectively). The variation of fluorescence properties allowed us to determine the dissociation constants (*K*_D_) for the complexes with the three UTRs G4, using excitation at 552 nm and emission recorded in the range 570-800 nm. The fluorescence spectra as well as the binding plots for UP1 and the three UTRs G4 are presented in **Figure 6**. In all three UTRs G4, the TAMRA fluorescence increased with the addition of UP1. This increase probably results from the Photoinduced Electron Transfer (PET) between the TAMRA fluorophore and the guanines of the G4s (46) This PET phenomenon leads to the TAMRA fluorescence quenching by the guanines of the G4, which is progressively minimized by the G4 disruption upon UP1 addition. The relative variations in the fluorescence intensities of UTR-Z and UTR-1 were smaller and were associated with lower affinities than UTR-C assuming 1:1 binding stoichiometry. Monitoring the maximum emission at approximately 580 nm, it showed a rapid saturation for UTR-C at 30-50 nM of UP1, a gradual plateau for UTR-1 at 50-100 nM, and a slow, extended response for UTR-Z at 500-1000 nM. The apparent dissociation constants (*K*_D_) were estimated as 8 nM for UTR-C, 20 nM for UTR-1, and 135 nM for UTR-Z, establishing the affinity order UTR-C > UTR-1 ≫ UTR-Z. These results follow the general trend observed by other methods were, indicating that UP1 binds most tightly to the UTR-C rG4 and least to both 1 and UTR-Z, the latter likely reflecting sequence polymorphism or conformational heterogeneity. Consistent with ITC and MST, fluorescence titration confirmed UTR-C as the highest-affinity UP1 substrate. In contrast, fluorescence measurements yielded an apparently weaker affinity for UTR-Z than for UTR-1, opposite to the trend observed by ITC and MST. This discrepancy reflects the fact that fluorescence titration in this setup reports not only binding occupancy but also the extent of G4 unfolding through photoinduced electron transfer (PET)-dependent TAMRA de-quenching. Although UP1 binds UTR-Z strongly, its pronounced conformational heterogeneity requires higher protein concentrations to fully disrupt all G4 conformers, leading to an increased apparent *K*_D_ in fluorescence titration.

**Figure 6.**
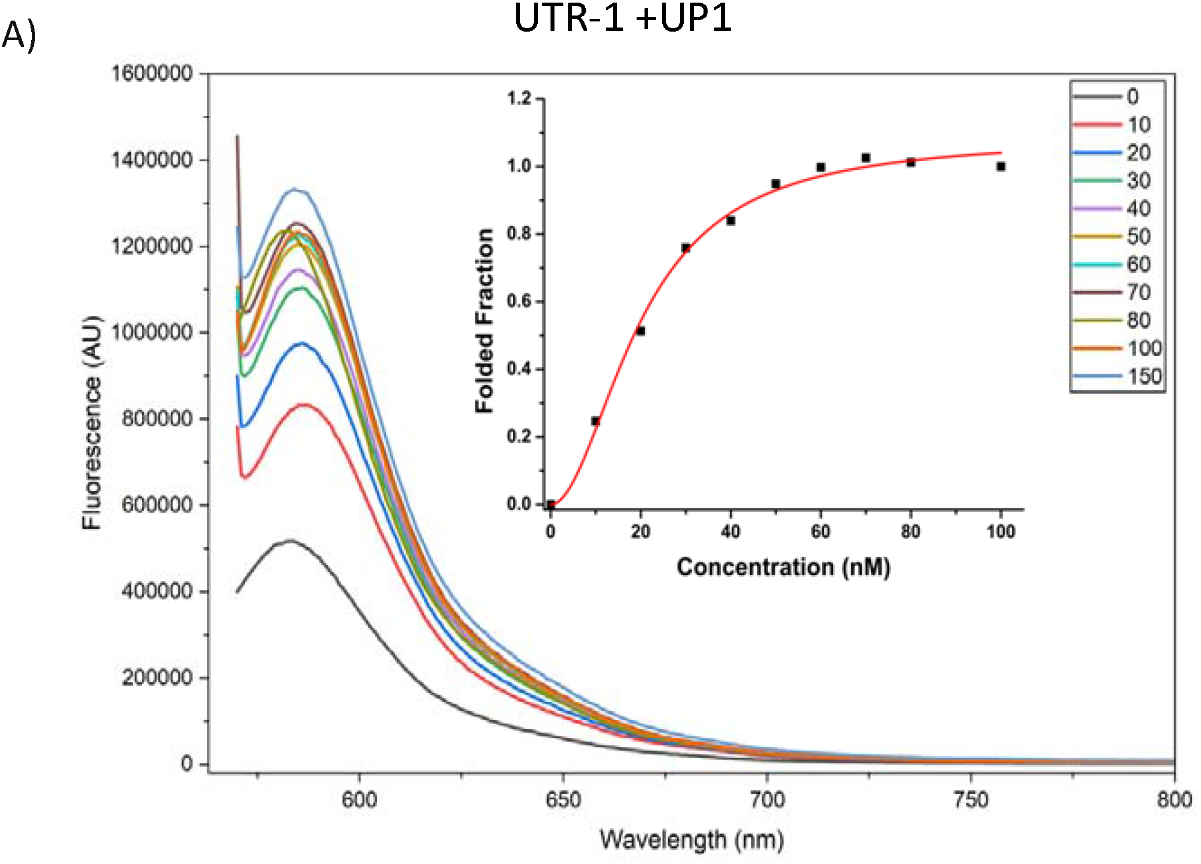
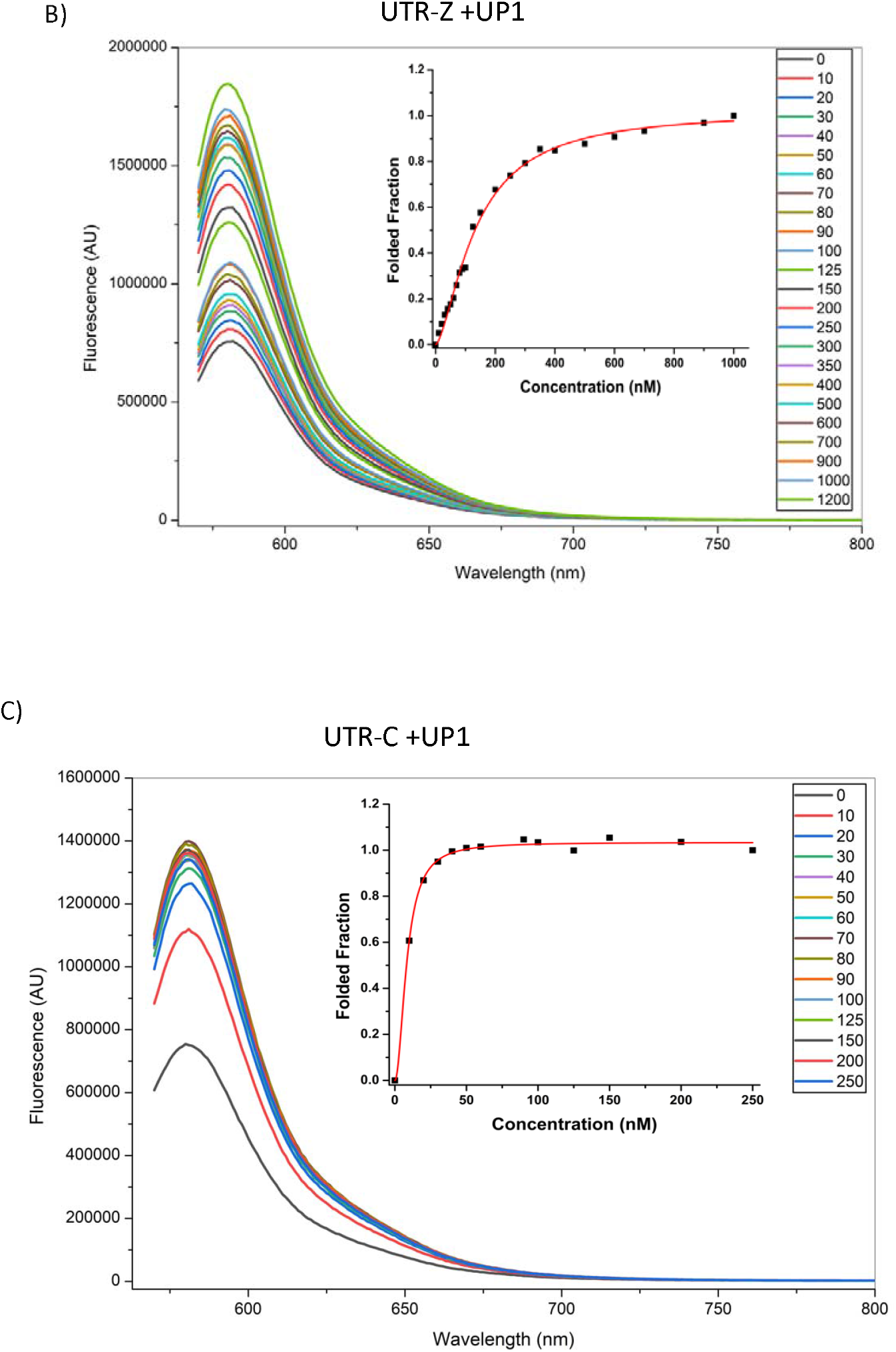
Fluorescence titration analysis of UP1 binding to double-labeled KRAS 5′UTR rG4s (A-C). Emission spectra (λ_ex_=552 nm) of 100 nM FAM/TAMRA-labeled rG4s upon UP1 addition. TAMRA de-quenching arises from disruption of guanine-photoinduced electron transfer (PET) upon G4 unfolding. UTR-C shows the tightest binding (KD ≈8 nM) and fastest saturation, reflecting optimal presentation of the canonical parallel fold; UTR-Z exhibits the weakest/slowest response, likely due to polymorphism reducing effective binding sites.

### NMR studies revealed the hnRNPA1 domains required for G4 recognition within the KRAS 5⍰UTR

To determine the molecular determinants of rG4 recognition by UP1, we performed 1D and 2D NMR experiments. In the presence of increased amounts of UP1, the typical imino proton peaks, characteristic of a folded UTR rG4 structure, disappeared proportionally to the amount of UP1 added to the sample. The results clearly indicate a complete disruption of the folded rG4 in complex with UP1 at molar ratios close to 1: 1 (**Figure 7A**). Additionally, 2D 1H-15N HSQC NMR experiments were performed using uniformly ^15^N-labeled UP1 to track changes in individual amino acids upon interaction with all three distinct RNA UTR G4s (UTR-Z, UTR-C, UTR-1). Three spectra are depicted in **Figure 7B**. Chemical shift deviations (Δδ/ppm) of amide groups in the most affected residues revealed structural changes upon rG4 binding and give hints about possible recognition motifs in the protein for each UTR rG4. UP1 contains two RNA-recognition motifs (RRM1 and RRM2), each with conserved RNP2 and RNP1 sub-motifs (14). Superimposed spectra before and after rG4 addition showed notable Δδ values for specific residues. Initially, the UP1 spectrum displayed well-separated and resolved peaks. Upon adding any of the UTRs rG4, the 2D ^1^H-^15^N HSQC NMR spectra depict two major types of peak changes: (i) Significant chemical shift variations (Δδ) and (ii) significant broadening and loss intensity of some peaks. In the first case, the Δδ results without significant peak volume reduction suggest binding modes compatible with conformational sampling induced by rG4 binding. These changes likely reflect a slight reorganization of the RRM1 residues near or within the principal binding site, causing indirect structural adjustments or subtle reorientations. To facilitate the analysis, we compiled all chemical shift perturbations (CSPs, ppm) per amino acid residue upon interaction of UP1 with each UTR G4 on Figure 8. The bar-plots represent CSP magnitudes for each residue across the protein sequence in presence of UTR-Z (green), UTR-C (blue) and UTR-1 (red). The horizontal orange dashed line indicates a threshold for significant CSPs, e.g., + 1 Standard Deviation (SD). In addition, we depict with triangles, the unambiguous residues with severe peak broadening or loss intensity on the superior part of Figure 8. These types of modifications are usually associated with intermediate exchange dynamics or tight binding at specific residues. To have a better perception of the binding mode and interaction sites, we depicted the residues with CSPs above 1 SD, as well as the residues with important peak loss or full disappearance on the UP1 structure (PDB: 2YIV; **Figure 9**. Residues with the largest Δδ values include S7, E12, Q13, L14, G21, E29, T40, T42, Y63, M73, H78, K79, K88, V84, V91, K162 and Q165; (ii) Intermediate exchange broadening for peaks, predominantly in the inter-RRM linker and RRM2 domain, suggests that RRM2 may serve as the primary rG4 binding domain. This effect likely arises from multiple binding modes and increased correlation time, leading to peak broadening or disappearance. Residues affected include S92, R93, E94, D95, S96, A101, L103, I108, L122, G130, M138, T139, D140, G142, G148, F151, F154, S159, K162, Y168, T170, N172, C176, R179, and A188. Many of these residues, especially F151 and F154, are located in the RNP1 motif of RRM2 (40). These observations collectively indicate that rG4s engage both RRM1 and RRM2, but with particularly strong effects observed in the RRM2 and in the inter-RRM linker. Three regions (residues ≈ 90-110, 138-146, and 160-182) emerged as common hotspots for all UTR rG4s. Interestingly, these coincide with functionally important regions: (a) the RRM1/RRM2 interface and linker, (b) the segment targeted by the small-molecule VPC-80051, which was designed to disrupt hnRNPA1 splicing activity in cancer (41), and (c) the RNA-binding groove where the short trinucleotide 5⍰-AGU-3⍰ was captured in the UP1-RNA crystal structure (42). Thus, our NMR data suggest that UTR rG4s exploit the same conserved RNA-interaction surface that is also vulnerable to small-molecule or oligonucleotide inhibitors. In line with these observations, aptamer BC15, an inhibitory RNA aptamer known to bind hnRNPA1, also targets overlapping RRM residues and competes with natural RNA substrates (43). BC15 likely stabilizes similar binding interfaces, thereby displacing rG4s and impairing UP1 function. Likewise, camptothecin, beyond its role as a topoisomerase I inhibitor, has been reported to interfere indirectly with hnRNPA1-RNA interactions, possibly by altering the same RRM2-linked conformational dynamics identified in our HSQC experiments (44). Both agents, like VPC-80051, converge on these sensitive RNA-binding regions, underscoring their relevance as therapeutic “Achilles’ heels” of UP1/hnRNPA1

**Figure 7.**
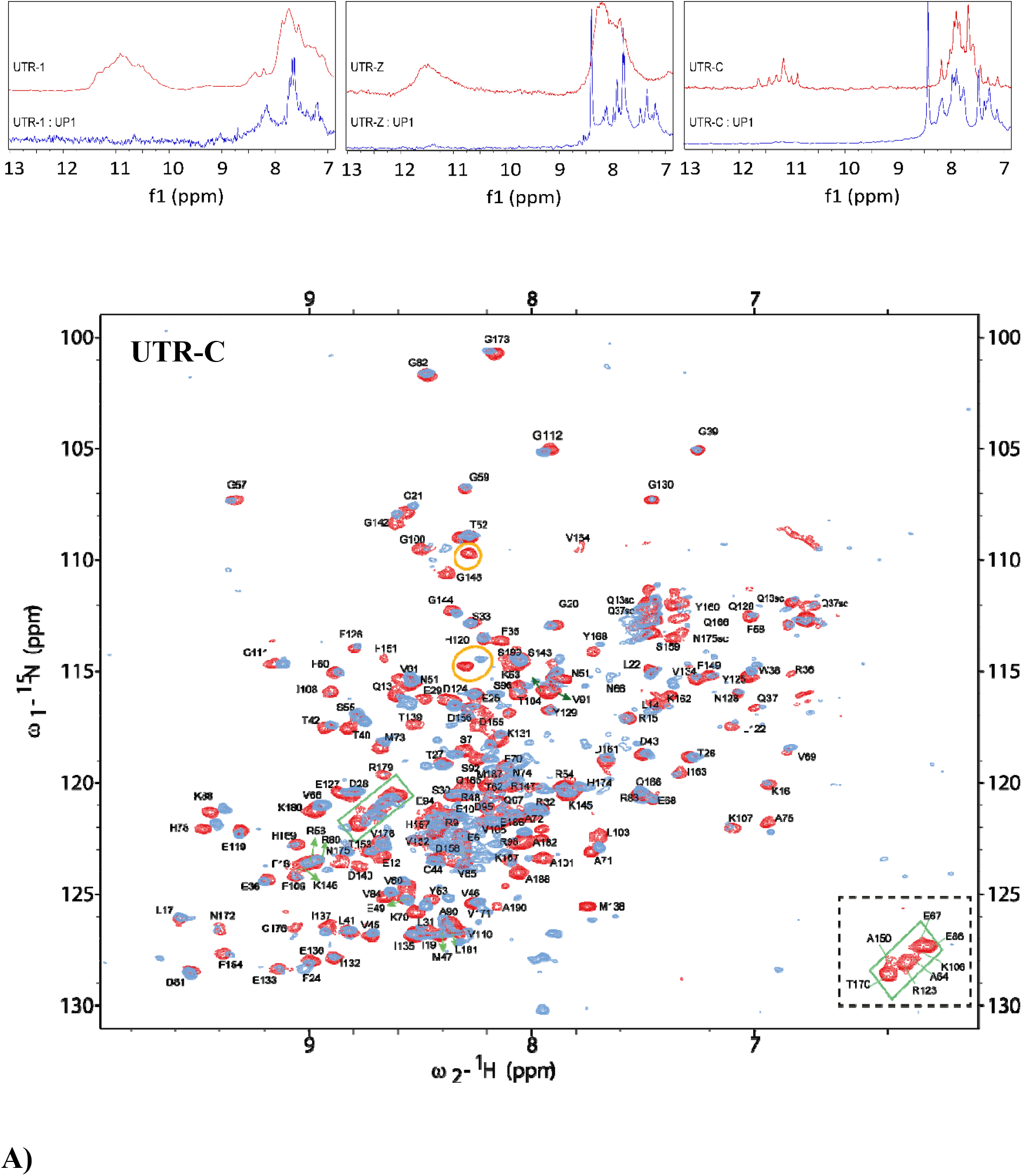
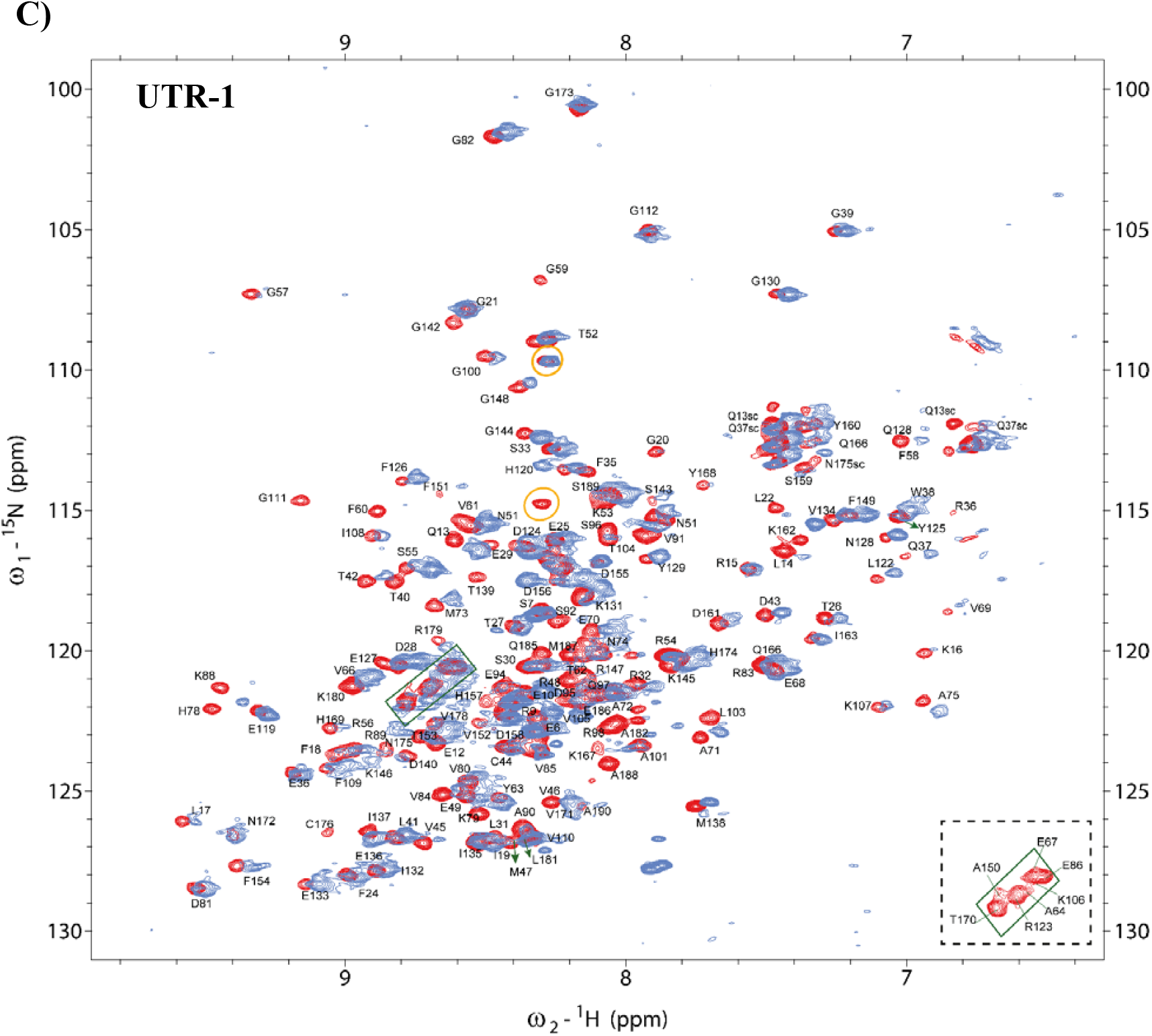
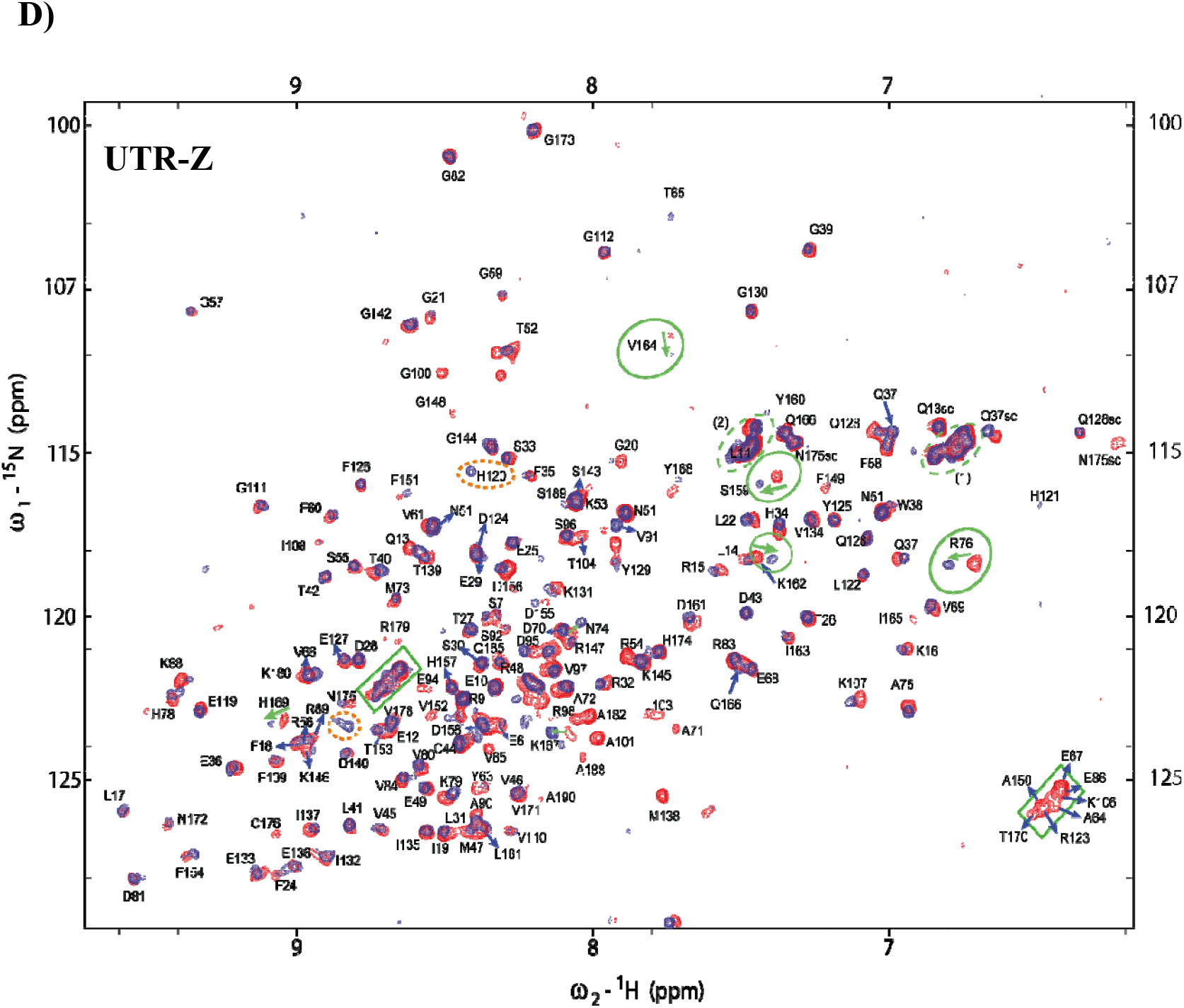
NMR characterization of UP1-rG4 interactions. 1D-NMR ^1^H spectra **(A)** depicting the imino and aromatic region of each UTR alone (red) and in presence of 1 molar equivalent of UP1 (blue), show near-complete loss of imino protons upon binding, indicating UP1-mediated disruption of Hoogsteen hydrogen bonds and G4 unfolding. **(B-D)** Overlaid ^1^H-^15^N HSQC spectra of ^15^N-UP1 (≈50 µM) before (red) and after (blue) addition of UTR-C (B), UTR-1 (C), or UTR-Z (D). Pronounced CSPs and severe line broadening/disappearance occur predominantly in RRM2, the inter-RRM linker, and parts of RRM1, suggesting that RRM2 serves as the primary binding interface, while the linker facilitates cooperative engagement of both RRMs during rG4 remodeling.

**Figure 8.**
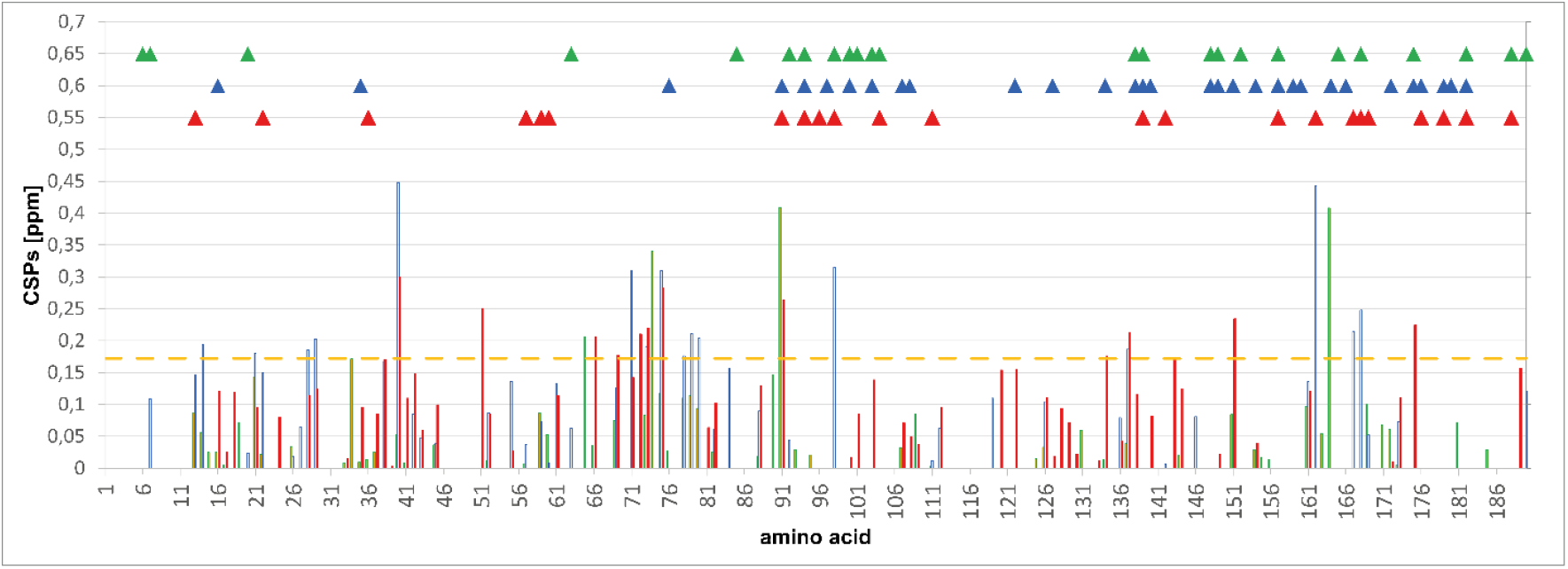
Chemical shift perturbation (CSP) analysis of UP1 upon binding to KRAS 5⍰UTR rG4s. The figure shows chemical shift perturbations (CSPs, ppm) per amino acid residue upon interaction of UP1 with three distinct RNA UTR G-quadruplexes (UTR-Z, UTR-C, UTR-1). The bar plots represent CSP magnitudes for each residue across the protein sequence. The horizontal orange dashed line indicates a threshold for significant CSPs (+ 1 Standard Deviation). Colored triangles above (green = UTR-Z, blue = UTR-C, red = UTR-1) mark residues where severe peak intensity losses or disappearance were observed in the HSQC spectra, often interpreted as strong binding, intermediate exchange, or conformational rearrangements.

**Figure 9.**
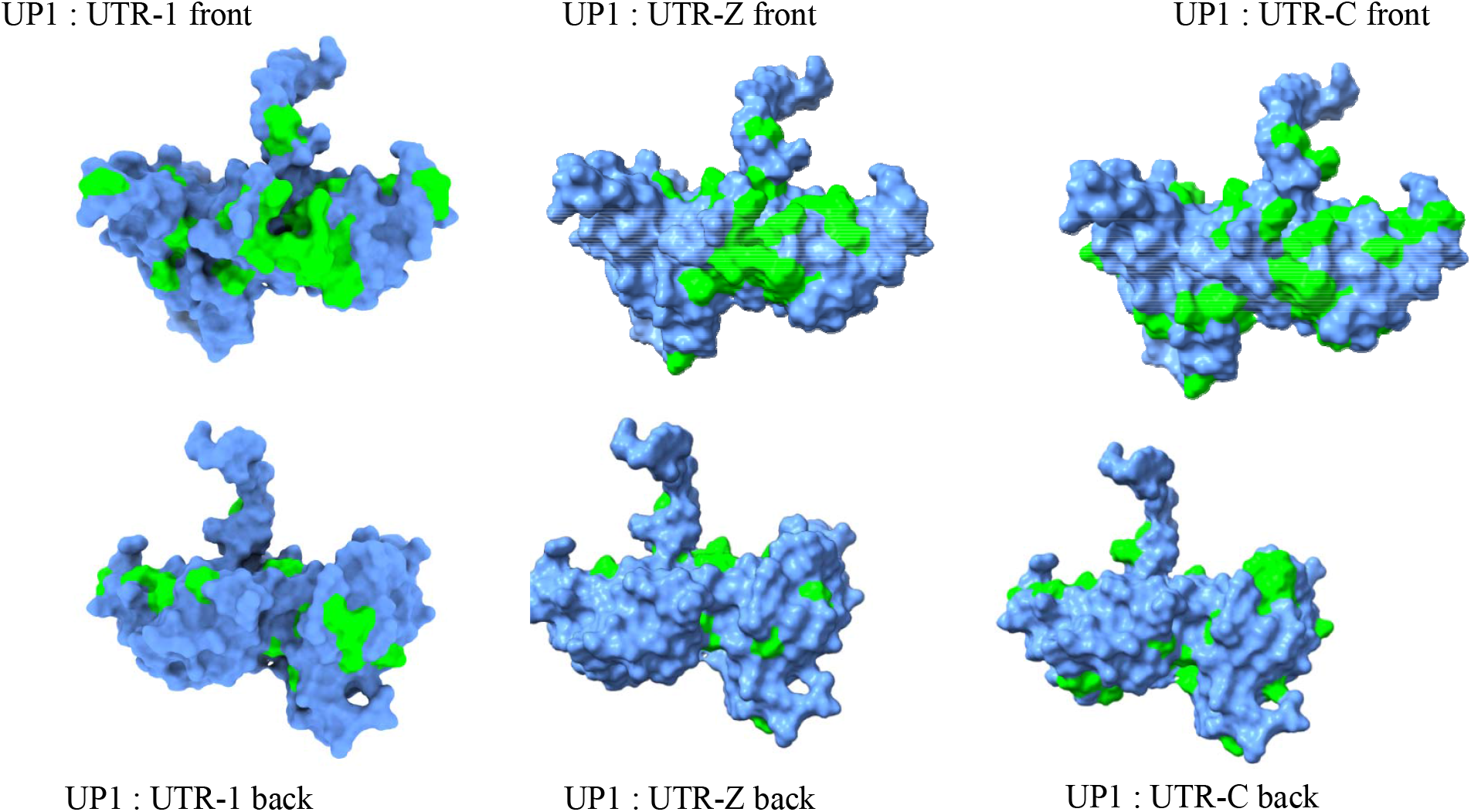
Mapping of rG4-sensitive residues onto the UP1 structure. Surface representations of UP1 (PDB: 2LYV) colored by CSP magnitude and severe broadening/disappearance (>1.5σ = green). Interaction hotspots concentrate on one face spanning both RRMs, with strongest effects in RRM2 and the interdomain linker. This unified binding surface enables UP1 to engage polymorphic rG4s through multivalent contacts, likely initiating unfolding by capturing transiently exposed single-stranded guanine tracts.

### Ligand stabilization and UP1 competition

The FRET-melting analysis of the three KRAS 5⍰UTR rG4 motifs (UTR Z, UTR 1, and UTR C) revealed clear differences in thermal stabilization in the presence of selected G4 ligands (**Figure 10**). The magnitude and direction of the Δ*T*_m_ values indicate that these oligonucleotides adopt distinct folding topologies and present variable ligand accessibility. These observations support the idea that the three motifs populate different conformational landscapes that accommodate ligands through distinct π-π stacking or groove-binding interactions, potentially also reflecting differences in their oligomerization state. These experiments were conducted with 360A, BRACO-19, and QUMA-1 at 1 and 2 molar equivalents.

**Figure 10.**
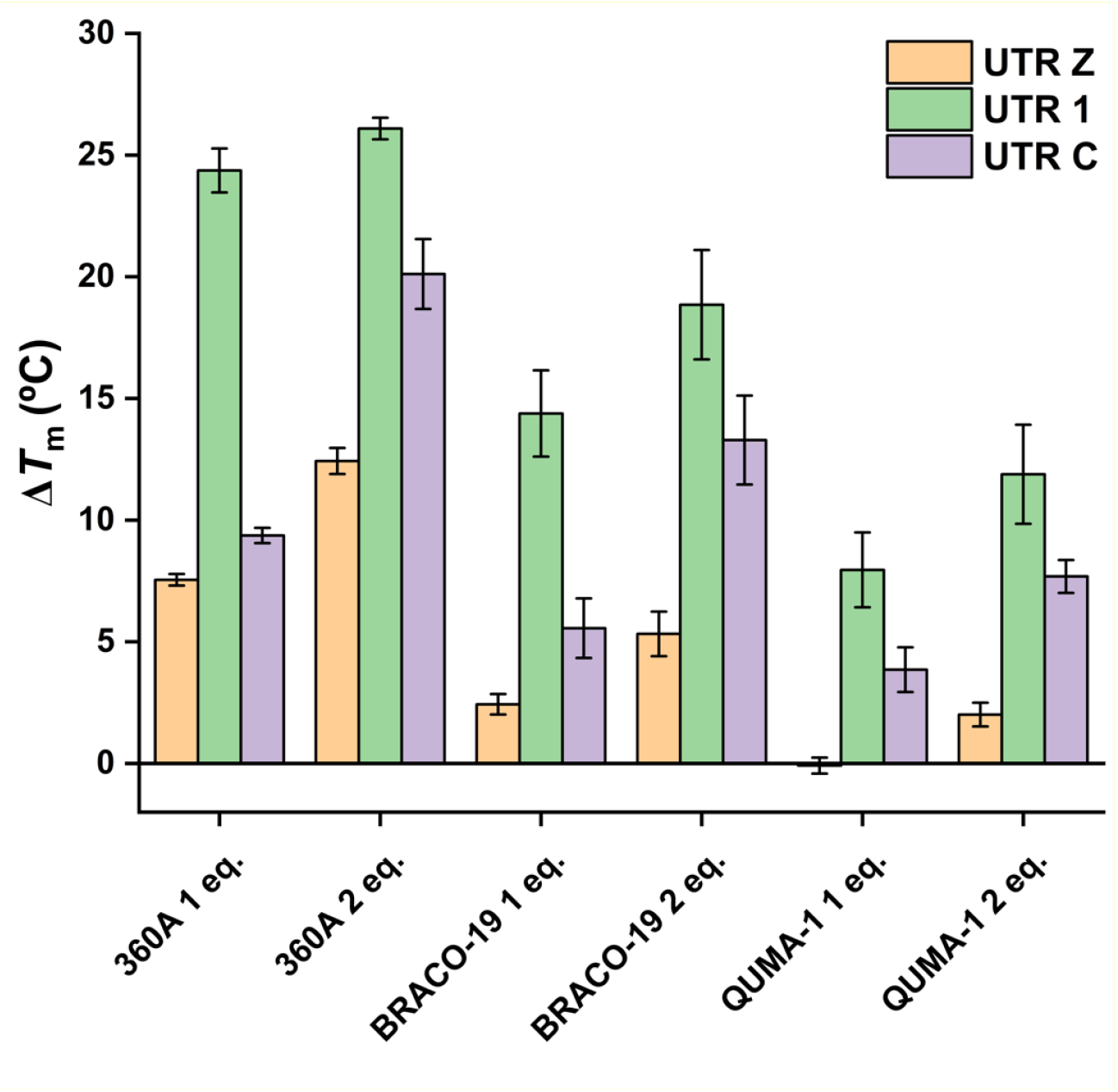
Ligand-induced thermal stabilization of KRAS 5UTR rG4s. Ligand-induced thermal stabilization (*ΔT*_m_, °C) of the tested sequences (0.2 μM) was evaluated in the presence of 1 or 2 molar equivalents of ligand. Experiments were performed in triplicate using 1×KPi buffer supplemented with 50 mM KCl (pH 6.7). Error bars represent the standard error of the mean (SE).

All ligands reproducibly induced thermal stabilization of the RNA sequences at 0.2 μM, with the exception of UTR Z in the presence of 1 molar equivalent of QUMA-1. Among the compounds tested, 360A produced the strongest effect. Notably, UTR 1 displayed the greatest stabilization across all compounds tested, and QUMA-1 the weakest one (**Figure 10**). PhenDC3 was also evaluated against the three sequences. Although the normalized melting curves were highly reproducible, they exhibited a characteristic profile with three plateaus, suggesting the presence of two melting events and likely reflecting the coexistence of multiple species. Because these curves could not be fitted reliably, no Tm values were extracted, and the corresponding data are shown separately (**Figure S1**).

To prove the robustness of these ligand-stabilized rG4s against an endogenous RNA-known unfolding factor, we employed fluorescence titrations with the UP1 protein (**Figure 11**). The fluorescence unfolding curves revealed that, despite prior stabilization by high-affinity ligands, UP1 efficiently remodeled the KRAS rG4s in a concentration-dependent manner. At sub-stoichiometric UP1 levels, the folded fraction decreased gradually, reflecting partial binding of the G4 structure by UP1. Once UP1 approached a 1:1 molar ratio with the RNA, a distinct transition occurred, evident from the steep inflection in the fluorescence signal around 100 nM of UP1, indicating a cooperative unfolding process leading to near-complete disruption of the G4 fold. The presence of this biphasic behavior suggests two mechanistically separable stages: an initial partial perturbation, likely involving transient binding to accessible loops or flanking regions non-occupied by the ligand, followed by full destabilization once UP1 saturates available binding sites. Accordingly, the current single-phase fitting model underestimates the complexity of this process, and a two-phase fitting model could better reflect the underlying kinetics and stoichiometry of the protein-RNA interaction before and after saturation. When comparing the two more stabilizing ligands, e.g., PhenDC3 and BRACO-19, both demonstrated substantial but incomplete protection against UP1-induced unfolding. The PhenDC3-stabilizing effect required slightly higher UP1 concentrations to achieve equivalent unfolding effects compared with BRACO-19, consistent with its higher thermal stabilization and possible tighter stacking at the terminal G-quartets. This differential “resistance” suggests that ligand-G4 complexes can transiently delay but not prevent protein-mediated remodeling. UP1, once sufficiently abundant, appears capable of opening the UTR G4 structures even in the presence of tightly bound stabilizers, presumably by binding to transiently exposed single-stranded guanine tracts and propagating local strand separation through cooperative RRM engagement. Such behavior implies that the energetic advantage of protein-RNA complex formation outweighs the ligand-induced stabilization energy, emphasizing UP1’s exceptional capacity to remodel structured RNAs. Mechanistically, these data reveal a competitive interplay between small-molecule stabilization and protein-mediated unfolding, with the outcome dictated by relative affinities, stoichiometry, and local structural dynamics. Ligands such as PhenDC3 are likely to reduce the rate of unfolding by restricting access of UP1 to the G4 core or by lowering the population of transiently unfolded intermediates. In contrast, UP1 exerts its effect through an entropic advantage of multivalent binding, as it was seen before (39, 45). Once one RRM domain engages a single-stranded region, it nucleates additional interactions that destabilize neighboring G-quartets. The biphasic unfolding kinetics thus capture the sequential transition from a ligand-protected to a protein-dominated state, where UP1 ultimately overrides chemical stabilization. This model aligns with prior reports of hnRNPA1 acting as a general G4 resolvase in mRNA 5⍰UTRs, promoting translation by eliminating structural impediments to ribosomal scanning. In addition, another mechanistic insight emerges from the incomplete fitting of the FRET curves: the two-phase profile indicates coexistence of subpopulations with distinct susceptibilities to UP1. One plausible interpretation is that ligand binding shifts the equilibrium toward a more homogeneous and thermally stable G4 topology, reducing conformational plasticity and delaying initial protein engagement. However, once the protein reaches stoichiometric saturation, the unfolding becomes cooperative and near-complete, implying that ligand protection is transient with an important dissociation rate *K*_off_. Such kinetic partitioning could represent a tunable regulatory mechanism in cells, where ligand-like metabolites or local protein concentrations modulate the folding equilibrium to fine-tune translation initiation efficiency.

**Figure 11.**
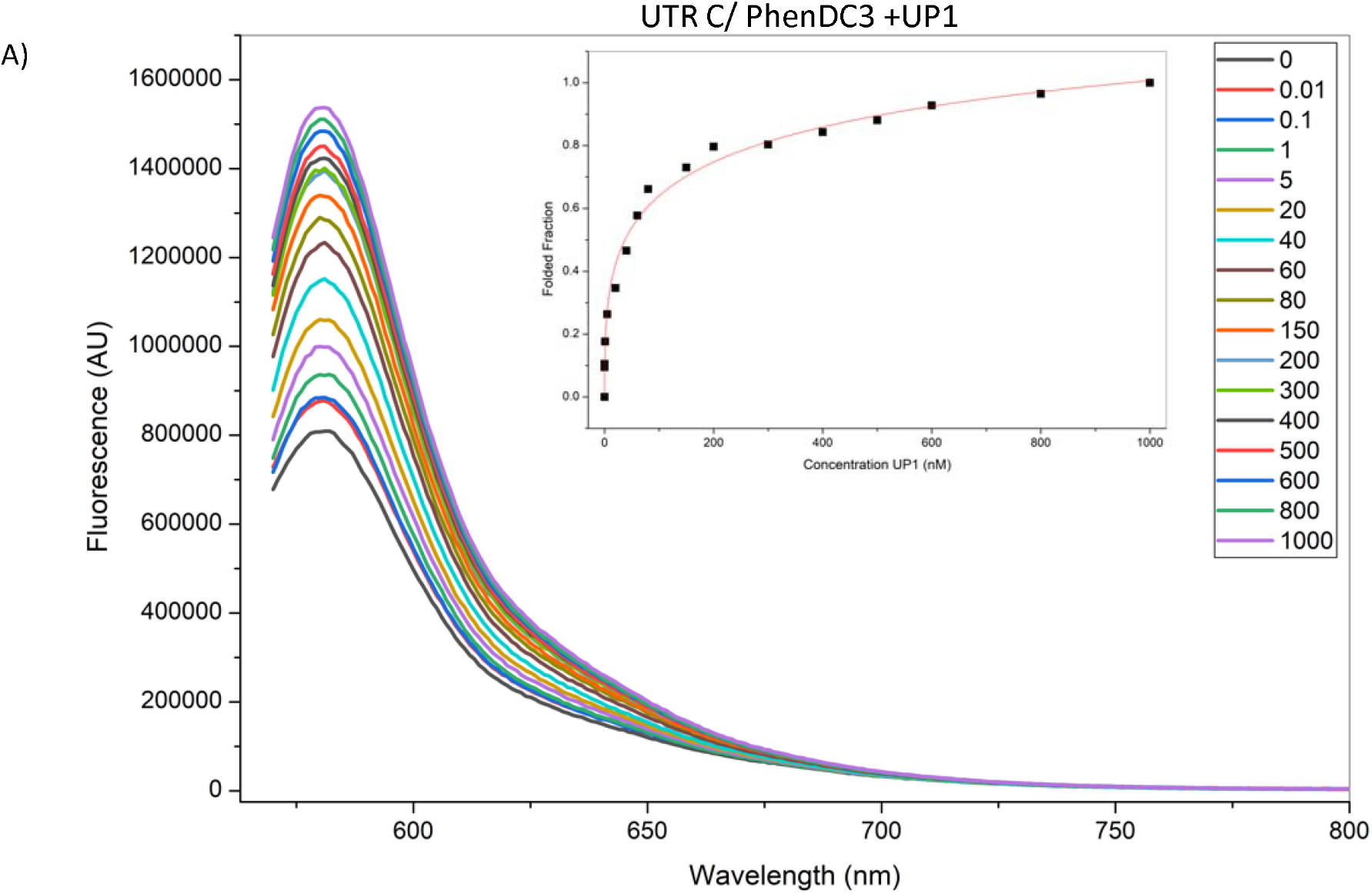
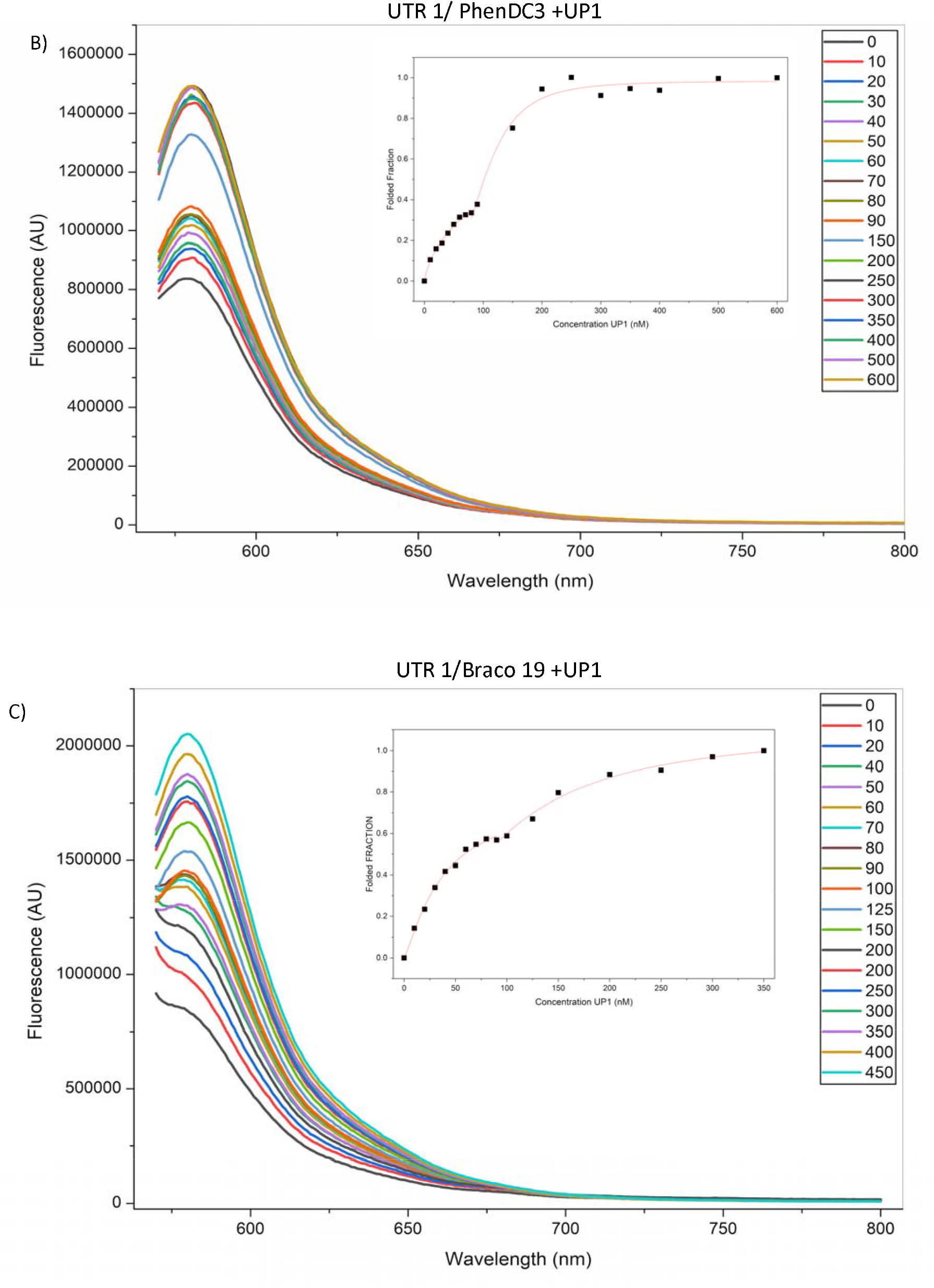
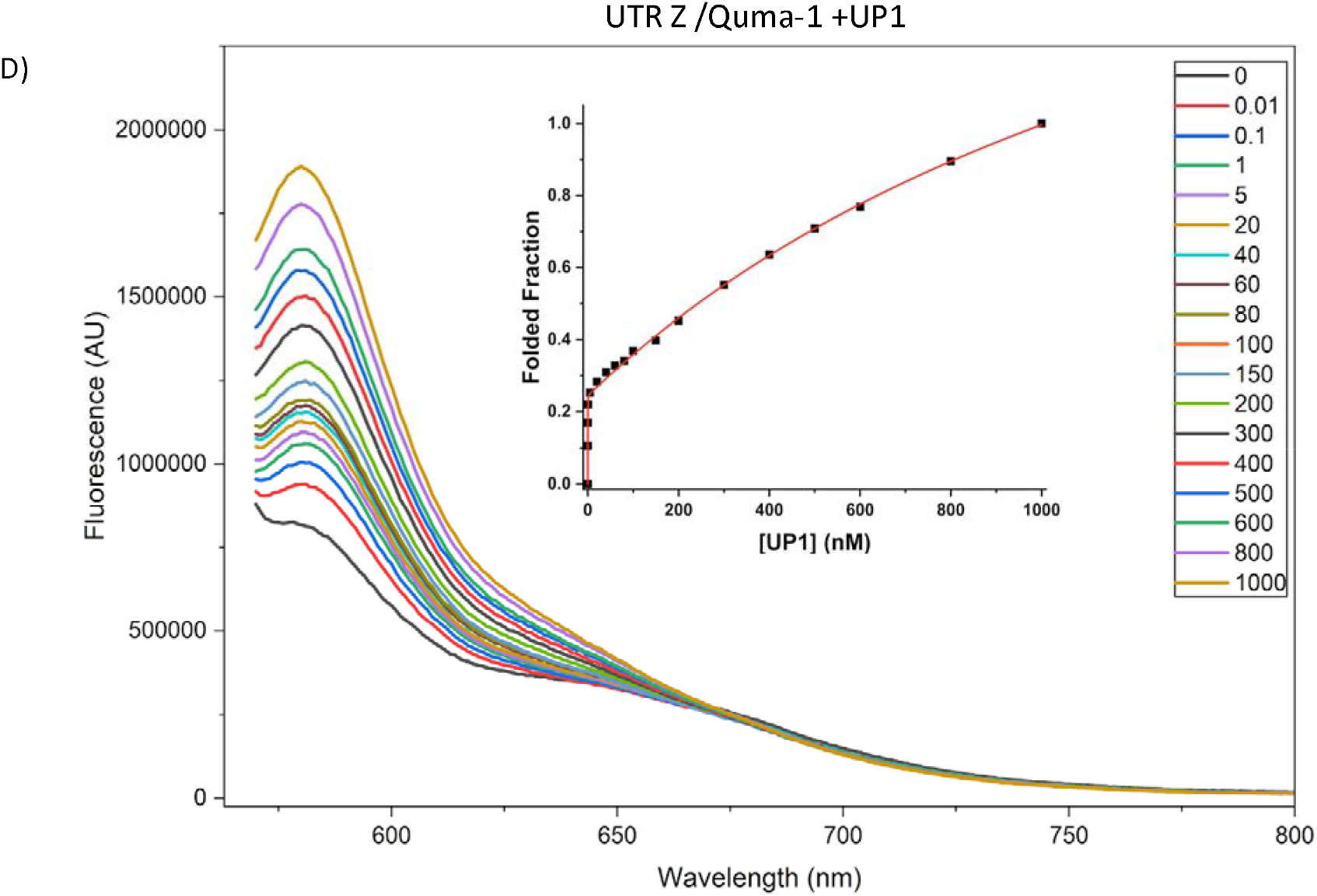
UP1-mediated unfolding of ligand-stabilized KRAS 5UTR rG4s. Fluorescence-based monitoring of UP1-induced unfolding of KRAS 5⍰UTR G-quadruplexes stabilized by ligands. These panels show fluorescence titration experiments monitoring the folded fraction of the UTR-C G-quadruplex in the presence of the stabilizing ligands PhenDC3 (A) and UTR-1 in PhenDC3 (B) and BRACO-19 (C) and UTR-Z in presence of Quma-1 in function of increasing UP1 protein concentration. Around 100 nM of UP1 (1:1 molar equivalent), a marked shift in the curves indicates a reduction in the folded population, suggesting that the protein effectively competes by overriding the ligand protection and emphasizing its role as a potent endogenous rG4 resolvase in KRAS translational control.

## Conclusion

Previous (38) cellular and biochemical studies demonstrated that hnRNPA1 binds KRAS 5⍰UTR rG4s, unfolds these structures, and promotes RNA remodeling and translational control. One of the main goals of this work was to use biophysical tools to decipher the molecular properties that govern the recognition and binding surfaces between UP1 and the G-quadruplexes present in the 5’ UTR mRNA of KRAS. In addition, we assessed whether this interaction can be perturbed by small molecules, potentially guiding future developments of selective ligands that target the RNA-protein complex. Although the three rG4 motifs differ markedly in stability, topology, and conformational dynamics, UP1 binds all of them through a conserved interaction surface spanning both RRMs with emphasis to RRM2 and the inter-RRM linker surface, as revealed by extensive chemical shift perturbations and exchange broadening in our NMR analyses. Notably, the binding affinities obtained across ITC, MST, and fluorescence titration converge to indicate that UP1 preferentially targets the most structurally homogeneous G-quadruplex, UTR-C, which adopts a well-defined, canonical parallel fold with minimal conformational exchange. In contrast, UTR-Z and especially UTR-1, which populate broader and more dynamic structural ensembles, exhibit weaker affinities despite their increased topological plasticity. These findings suggest that UP1 does not merely select for conformational heterogeneity but instead recognizes specific architectural features that are optimally presented in a well-folded rG4 scaffold. Mechanistically, UP1 acts as a potent rG4 resolvase even highly stabilized rG4-ligand complexes, such as those formed with PhenDC3 or BRACO-19, are ultimately remodeled by UP1 once stoichiometric saturation is reached. This competitive behavior underscores a regulatory hierarchy in which small-molecule stabilization can transiently modulate the folding landscape, yet protein-mediated engagement and unfolding dominate the equilibrium. Such a mechanism provides a structural basis for how hnRNPA1 facilitates KRAS translation by resolving rG4 barriers that impede ribosomal scanning within the 5⍰UTR. In addition, we show that deeper and more structurally-oriented studies are necessary to understand how ligands could be improved to compete better against proteins such as hnRNPA1. Our binding and structural analyses refine the model proposed in previous studies (38). In that work, UTR-Z and UTR-C were identified as the rG4 elements that are most likely to form stably in the cellular context, whereas UTR-1 remains partially unfolded. Here, we show that among these folded elements, UP1 exhibits the highest affinity for the structurally homogeneous and canonical UTR-C rG4, while UTR-Z and UTR-1 bind progressively weaker in accordance with their increased conformational heterogeneity. Another result that indicates a rather structural recognition by UP1 (the T_M_ values), than just a folding equilibrium, is the fact that UTR-C and UTR-Z represent the least and the most thermodynamically stable structures respectively. By establishing the molecular logic through which UP1 discriminates among distinct rG4 topologies and remodels them, this study provides a foundation for designing RNA-targeted ligands able to stabilize specific KRAS rG4s against endogenous G4-resolving factors. Such strategies may enable selective attenuation of KRAS translation in cancers driven by its dysregulated expression.

## Materials and Methods

### Sequences used in this work

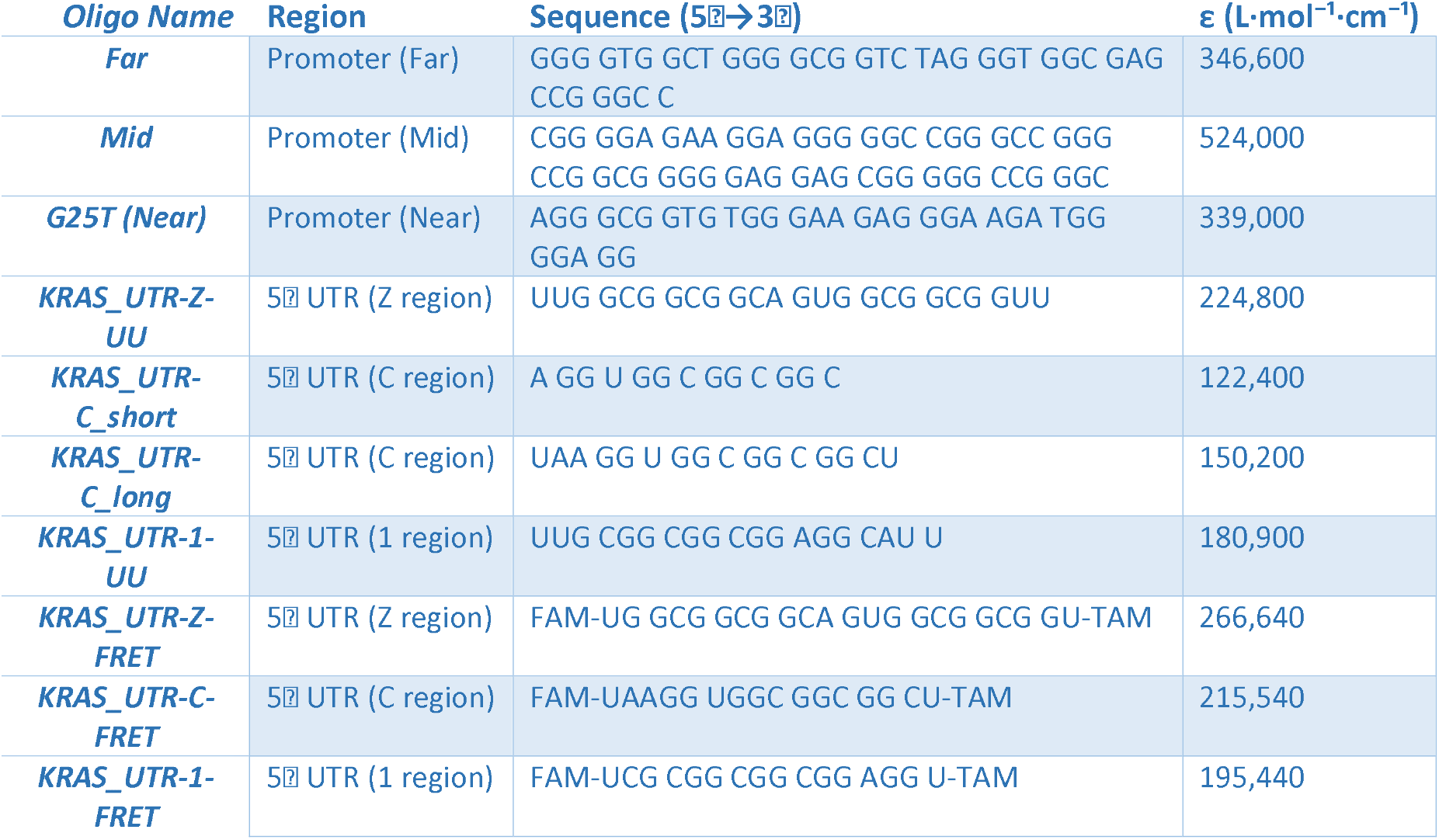

#### Heat-shock transformation of E. coli DH5α (pGEX-6P-1)

15 µL of chemically competent Escherichia coli DH5α were mixed gently on ice with 10 ng of the pGEX-6P-1 plasmid (negative controls received the same volume of ddH_2_O instead of DNA). Cell-DNA mixtures were incubated on ice (4 °C) for 30 min, heat-shocked at 42 °C for 1 min, and immediately returned to ice for 5 min. For recovery, 900 µL of antibiotic-free LB broth was added and cultures were shaken at 37 °C for 30-60 min. Cells were then spread onto LB-agar plates supplemented with ampicillin and incubated overnight at 37 °C. Supplementary figure S2.

### Post-transformation screening and culture

#### Bacterial cultures

To validate the transformation, three plates were prepared: a viability control consisting of competent cells plated on LB without antibiotic (expected growth), an antibiotic control consisting of competent cells plated on LB + ampicillin (expected no growth), and the transformation sample consisting of the recovered cells mixed with pGEX-6P-1 plated on LB + ampicillin (expected distinct colonies). The following day, well-isolated colonies from selective plates were picked individually and grown overnight at 37 °C with shaking in 5 mL LB containing ampicillin for downstream analyses.

#### Plasmid extraction (Miniprep)

Plasmid DNA was isolated from 2-5 mL overnight LB-ampicillin cultures using a silica spin-column plasmid miniprep kit (e.g., QIAprep Spin Miniprep, Qiagen) according to the manufacturer’s instructions. To verify plasmid topology and rule out nicking, 200-500 ng of each miniprep was analysed on a 1.0% agarose gel in 1× TAE at 5-7 V·cm^−1^ for 45-60 min alongside a supercoiled DNA ladder and the same plasmid linearised with a single-cut restriction enzyme.

Gels were stained with ethidium bromide (0.5 µg·mL^−1^) or GelRed and imaged under UV; under these conditions the supercoiled form migrates fastest, the linear form runs at the size-expected position, and the open circular (nicked) form migrates more slowly. Where greater resolution of topoisomers was needed, an aliquot was relaxed with E. coli Topoisomerase I (37 °C, 15-30 min) and/or separated on chloroquine-agarose (2-5 µg·mL^−1^ chloroquine).

### Expression and Purification of the UP-1 Fragment of HNRNPA1

The UP-1 fragment of HNRNPA1 was expressed from the transformed DH5α as mentioned before a GST fusion protein using the pGEX vector system. Expression was induced in E. coli cultured in Terrific Broth (TB) medium supplemented with phosphate buffer. When the culture reached an OD_600_ between 1.5 and 2.0, protein production was induced by the addition of 1 mM IPTG. Cultures were incubated overnight at 17⍰°C. Cells were harvested by centrifugation at 5,000 × g for 20 minutes at 4⍰°C. Bacterial pellets were ressuspended in 1× PBS containing 0.02 mg/mL lysozyme and 1 mM PMSF. The suspension was incubated for 30 minutes at 4⍰°C with gentle stirring. Cell lysis was performed by sonication at 3 W, using three cycles of 30 seconds each, separated by one-minute intervals on ice. After the second cycle, a brief synchronization step (sync) was introduced prior to the final sonication to improve consistency and minimize foam formation or overheating. The lysate was clarified by ultracentrifugation at 20,000 rpm for 1 hour at 4⍰°C. The resulting supernatant was applied to a GSTrap affinity column pre-equilibrated with 1× PBS. The column was washed with 1× PBS containing 1 M NaCl, and the fusion protein was eluted using 20 mM reduced glutathione in 1× PBS (pH 8.0). The GST tag was removed by incubation with PreScission protease, followed by dialysis against 1× PBS containing 1 mM DTT. The cleaved UP-1 fragment was further purified by incubation with glutathione agarose beads equilibrated in 1× PBS to remove any residual GST or uncleaved fusion protein. Protein purity was assessed by SDS-PAGE, and concentration was determined spectrophotometrically using the predicted extinction coefficient.

### Generation of Panc-1 Cells with KRAS 5⍰UTR Reporter Activity

The KRAS 5⍰UTR fragments (utr-1, utr-c, utr-z) were cloned by oligo cloning into the lentiviral vector pLV-mScarlet-Control-5⍰UTR (Addgene, USA) digested with Esp3I and AgeI. HEK293 cells were transfected with 10 μg of each lentiviral construct separately, along with 1.8 μg of pMD2.G and 5 μg of pPAX2. After 48 hours, viral particles were collected, filtered, and used to infect Panc-1 cells, either wild-type or HNRNPA1 knockout, previously characterized (38).

### RNA-Immunoprecipitation (RIP)

Panc-1 cells (30 × 10^6^) were collected and lysed for 10 min in hypotonic lysis buffer (5 mM Pipes, 85 mM KCl, 0.5% NP-40) supplemented with a protease inhibitor cocktail. Nuclei were removed by centrifugation, and the cytoplasmic fraction was diluted 1:1 with G4RP buffer [150 mM KCl, 25 mM Tris (pH 7.4), 5 mM EDTA, 0.5 mM DTT, and 0.5% NP-40 in nuclease-free water].

After pre-clearing, lysates were incubated with 3 μg anti-hnRNPA1 antibody (Merck, clone 9H10) or IgG control, together with 8 μl Dynabeads Protein G (Thermo Fisher Scientific, USA) for 6 h at 4°C with gentle rotation. Beads were washed thoroughly and RNA was extracted using 1 ml Tri-Reagent (Sigma-Aldrich, Germany) following a phenol/chloroform protocol. RNA was treated with DNase I (Thermo Fisher Scientific) to remove genomic DNA.

RNA concentration, integrity, and purity were assessed using the Agilent 2100 Bioanalyzer (RNA 6000 Nano Kit). 100 ng of RNA was reverse transcribed into cDNA using the High-Capacity RT Kit (Thermo Fisher Scientific, USA). Quantitative PCR (qPCR) was performed using SYBR Green chemistry (KAPA Biosystems, USA) with the following primers:

- Forward: 5⍰-AAGCTGAAGGTGACCAAGGG-3⍰
- Reverse: 5⍰-GCCGTCCTCGAAGTTCATCA-3⍰

Expression levels were normalized to IgG control and quantified using the ΔΔCt method.

### Circular Dichroism (CD) Spectroscopy

CD spectroscopy was performed to assess the secondary structures of the three RNA constructs corresponding to the 5⍰ untranslated regions: UTR-Z, UTR-C, and UTR-1. RNA samples were prepared at a final concentration of 1 µM in 10 mM potassium phosphate buffer (pH 7) containing 50 mM KCl. Prior to measurements, the RNAs were heated to 95°C for 5 minutes and slowly cooled to room temperature to promote proper folding. CD spectra were recorded using a JASCO J-600 spectropolarimeter equipped with a thermostatted cell holder and a 1 cm pathlength quartz cuvette. A thermometer placed in the cuvette holder ensured accurate temperature monitoring. Spectra were acquired in the 220-340 nm range, averaged over three scans, smoothed, and baseline-corrected by subtracting the buffer signal. Data were reported as ellipticity (mdeg) versus wavelength and analyzed using J-700 Standard Analysis Software (Japan Spectroscopic Co., Ltd).

### Thermal Denaturation Experiments

Thermal denaturation profiles were obtained by monitoring the CD signal at 263 nm while gradually increasing the temperature from 5°C to 95°C. The measurements were performed using the same buffer conditions as described above. Data were collected at 1°C intervals, with a 1-minute equilibration at each temperature point. Melting curves were generated by plotting ellipticity as a function of temperature, and the first derivative of the CD signal (dθ/dT) was calculated to determine the temperature corresponding to the structural transition. All experiments were conducted in triplicate under identical conditions.

### CD Titration experiment for promoter

CD spectra have been obtained with a JASCO J-600 spectropolarimeter equipped with a thermostatted cell holder.CD experiments were carried out with the three oligonucleotides near mid far (3 µM) in 50 mM K-phosphate buffer, pH 7, 50 mM KCl alone and with and increasing amount of UP1. Spectra were recorded in 1 cm quartz cuvette. A thermometer inserted in the cuvette holder allowed a precise measurement of the sample temperature. The spectra were calculated with J-700 Standard Analysis software (Japan Spectroscopic Co., Ltd) and are reported as ellipticity (mdeg) versus wavelength (nm). Each spectrum was recorded three times, smoothed and subtracted to the baseline.

### Förster Resonance Energy Transfer (FRET) Melting Experiments

FRET-melting experiments were performed on the CFX Connect Real-Time PCR Detection System (Bio-Rad, Hercules, CA, USA), equipped with a FAM filter (λ_ex_=492 nm; λ_em_=516 nm), and oligonucleotides used were labeled with fluorescein (FAM) and rhodamine (TAMRA) at the 5’ and 3’ ends, respectively. Before the experiment, oligonucleotides at 0.25 μM were annealed in 1x Kpi buffer as detailed in the preceding sections. Then, 20 μL of oligonucleotides were aliquoted into each strip for each condition, followed by 5 μL of ligand from (100 μM stock), to reach a final concentration of 5 molar equivalents. For control conditions, 1x KPi buffer was added. The thermocycler was parameterized to measure and acquire the FAM emission after each step with a gradual increase of 0.5⍰°C every 1 min, from 25⍰°C to 95⍰°C. The normalized fluorescence curves were used to estimate the melting temperatures. The *T*_m_ values were emission to 0.5. For each sequence and ligand, three independent experiments were carried out.

### Fluorescence titrations

Fluorescence titrations were performed on a HORIBA FluoroMax-4 fluorometer equipped with a Peltier temperature controller. FRET-modified RNA sequences (UTR-Z, UTR-C, and UTR-1) bearing 5⍰-6-FAM and 3⍰-TAMRA-Sp were annealed at 100 nM in 700 µL 1× KPi buffer containing 50 mM KCl and transferred to a quartz Suprasil cuvette (10 × 4 mm; Hellma, ref. 104F-10-40). UP1 protein from a 58 µM stock was added directly to the cuvette in small aliquots to achieve increasing protein concentrations while maintaining a constant RNA concentration. After each Spectra, samples were incubated for 3 min and emission spectra (570-800 nm) were recorded as the average of three scans (integration time 0.5 s). Excitation/emission slit widths were, respectively: UTR-Z, 5/4 nm; UTR-C, 6/7 nm; UTR-1, 6/5 nm. Measurements were carried out at 25 °C.

For each titration point, the fraction of UP1-bound RNA (α) was calculated from the fluorescence intensity at 580 nm using Eq. (1), where I is the observed intensity, I^free^ the intensity of RNA alone, and I^bound^ the intensity at saturating UP1:

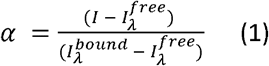

Data points were then fitted into the Hill saturation binding model using OriginPro 2021, and apparent equilibrium constants (*K*_D_). In which *K*_D_ is the apparent equilibrium dissociation constant, [RNA] is the concentration of the RNA and *n* is the Hill constant which defines the cooperativity of UP1 binding.

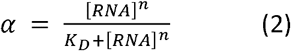

### Ligand competition fluorescence titration

Ligand competition experiments were carried out by fluorescence titration, using the same instrument, buffer, cuvette, acquisition settings, and analysis pipeline described above. RNA ligand mixtures (1:1 molar ratio; 100 nM RNA in 1× KPi, except Quma-1 was at 1:2 RNA/ligand ratio) were prepared by pre-equilibrating the FRET-labeled RNA (5⍰-6-FAM / 3⍰-TAMRA-Sp) with ligand at room temperature, then transferred to the cuvette. UP1 from the same stock solution of 58 µM was added in small aliquots to increase UP1 concentration while keeping RNA and ligand concentrations constant. After each addition, spectra (570-800 nm) were recorded as above.

### Binding Analysis by ITC

Isothermal titration calorimetry (ITC) experiments were carried out using a MicroCal VP-ITC instrument to determine the thermodynamic parameters of interaction between UP1 and different G-quadruplex-forming sequences from the KRAS promoter. To preserve the native G4 structure, all oligonucleotides were pre-folded and diluted in 1× potassium phosphate buffer (KPi), a condition known to stabilize G-quadruplexes. The exact buffer composition was 10 mM potassium phosphate, pH 7.4, prepared with nuclease-free water. Due to the limited yield of UP1 protein, reverse titration was performed, with 8⍰μM UP1 placed in the sample cell and G4 oligonucleotides loaded into the injection syringe. Three DNA constructs were tested, representing G4 sequences located at near, mid, and far positions relative to the canonical 32R region of the KRAS promoter. For the near and mid sequences, the oligonucleotide concentration in the syringe was 80⍰μM. For the far sequence, 60⍰μM was used to prevent early saturation. Each titration consisted of 2⍰μL injections, performed over 4 seconds, with 200 seconds spacing between injections. All experiments were carried out at 25⍰°C under identical buffer and concentration conditions to ensure comparability. Data were analyzed using a single-site binding model in the MicroCal Origin software.

### Microscale Thermophoresis (MST)

MST experiments were conducted in triplicate using a Monolith NT.115 system (NanoTemper Technologies). Cy5-labeled RNA oligonucleotides (UTR-1, UTR-C, and UTR-Z) were used as ligands, and the UP-1 protein was used as the binding target. All oligonucleotides were prepared in MST buffer containing 1× KP buffer supplemented with 0.05% Tween-20 to minimize nonspecific interactions. Each Cy5-labeled oligonucleotide was used at a fixed final concentration of 5 nM in the binding reactions. A 16-point, 2-fold serial dilution of UP-1 protein was prepared in the same buffer, starting at 100 µM in tube 1 and ending at approximately 1.5 nM in tube 16. For this, 20 µL of UP-1 solution at the highest concentration was added to tube 1, and 10 µL of buffer was added to tubes 2 through 16. Serial dilutions were performed by transferring 10 µL from one tube to the next, discarding 10 µL from tube 16 to maintain equal volume. Subsequently, 10 µL of Cy5-labeled oligonucleotide was added to each tube in reverse order (from tube 16 to tube 1) to ensure consistent incubation time. Each UTR oligo was tested in an independent experiment. After a brief incubation at room temperature, samples were loaded into standard capillaries (NanoTemper Technologies). MST measurements were carried out using 20% LED power and 40% MST power. Data analysis was performed using the NanoTemper Analysis software. Normalized binding curves were generated using the temperature-jump (T-Jump) method and fit to a single-site binding

## Acknowledgements

Zahraa Othman is recipient of the Excellence Eiffel fellowship from France’s Ministry of Europe and Foreign Affairs and the University of Lebanon. Pedro Lourenço acknowledges a fellowship grant from CICS-UBI (ref. CICS UBI_BI_LABORATÓRIOS DE INVESTIGACAO_2025) and a doctoral fellowship grant from FCT (ref. 2025.02136.BD). David Moreira acknowledges a doctoral fellowship grant from FCT, reference (https://doi.org/10.54499/2023.02001.BD). We also would like to acknowledge Campus France for the Programme PESSOA (ref. 52857XG) support. With financial support from the 2021-2030 Cancer Control Strategy (PCSI), on funds administered by Inserm (ref. ASC24034GSA).

## Supplemental Information

**Supplementary figure 1.**
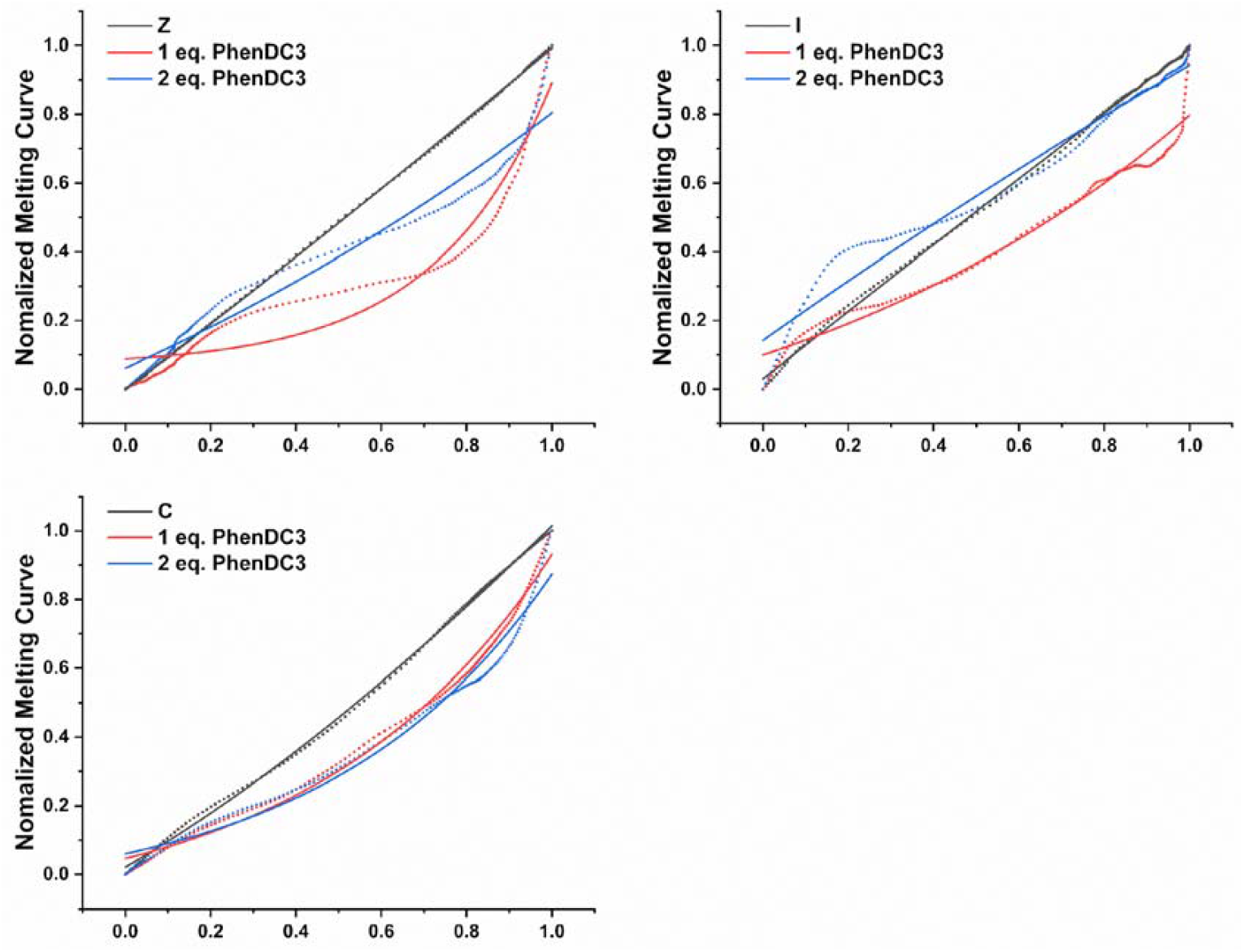
PhenDC3 results for the tested sequences. The data represent the mean of normalized values, plotted after analysis. Interestingly, the melting curves display a similar profile across all sequences, showing three distinct plateaus that suggest a dual thermal stabilization effect, likely corresponding to the melting of two species. These results are not included in the previous bar graph, as a reliable curve fitting could not be achieved, and therefore, no melting temperature (Tm) values were determined.

**Supplementary Figure 2.**
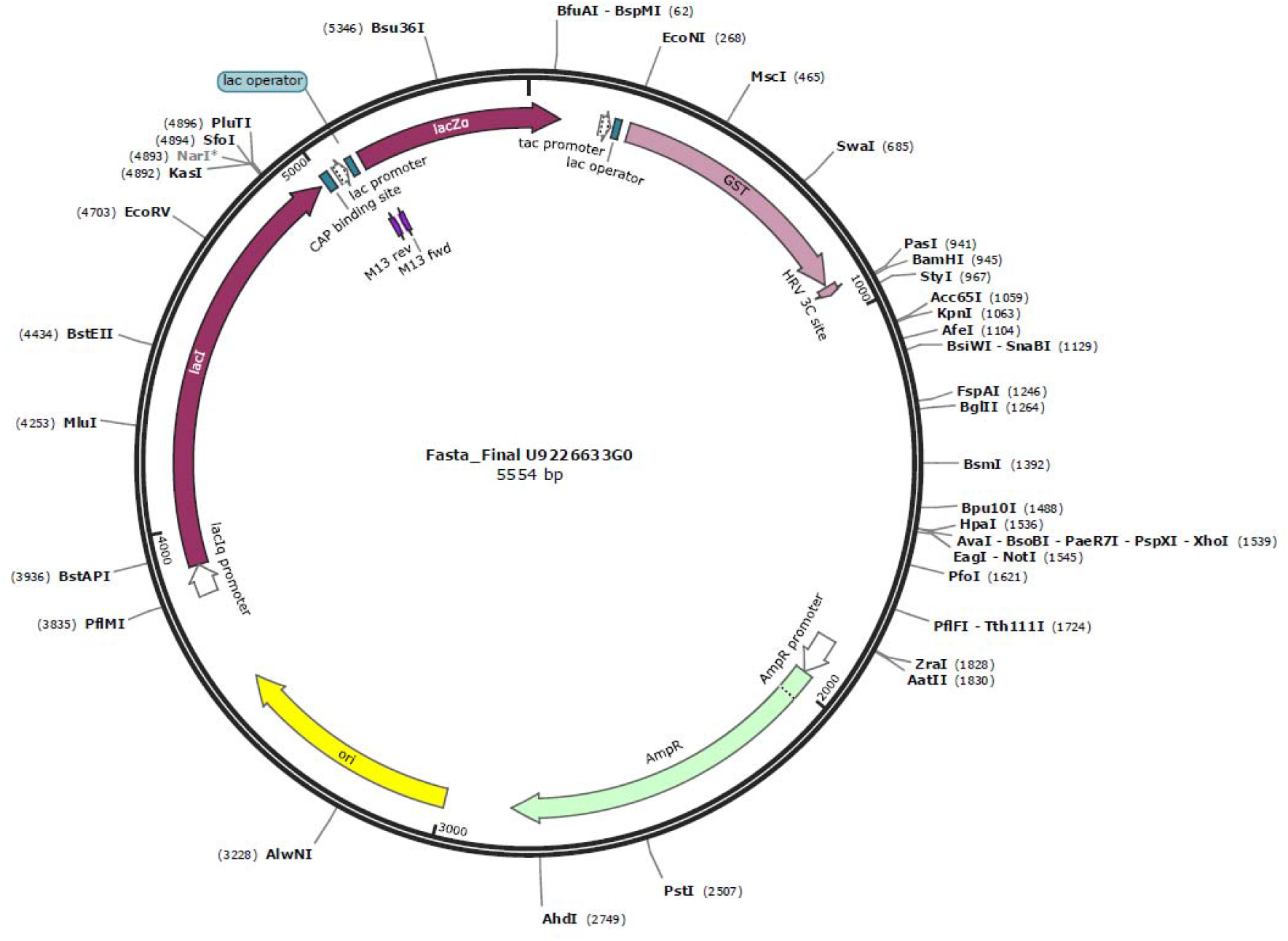
Plasmid used in transformation to express UP1 contains ampicillin resistance gene.

